# Improving positively tuned voltage indicators for faster kinetics and higher contrast

**DOI:** 10.1101/2024.06.21.599617

**Authors:** Sungmoo Lee, Guofeng Zhang, Yukun Alex Hao, Richard H. Roth, Yue Sun, Can Dong, Jun Zhu, Laura C. Gomez, Ryan G. Natan, Dongyun Jiang, Lan Xiang Liu, Guilherme Testa-Silva, Atsuki Hiramoto, Botond Roska, Daniel Feldman, Na Ji, Thomas R. Clandinin, Lisa Giocomo, Jun Ding, Michael Z. Lin

## Abstract

Existing positively tuned ASAP-family voltage indicators such as ASAP4e exhibit superior photostability compared to the negatively tuned ASAP3 and ASAP5, but signal-to-noise ratios for spike detection remained similar to ASAP3. To improve spike detection by positively tuned ASAP indicators, we performed multiple rounds of structure-guided saturation mutagenesis of an ASAP4e predecessor while screening directly for faster responses. One variant, ASAP6c, demonstrated ∼100% increases in fluorescence in response to single action potentials in both one-photon and two-photon imaging in vivo, achieving a 3-fold improvement in signal-to-noise ratios over ASAP4e.

---

An overarching goal of neuroscience is to understand how circuits of neurons represent, store, and transmit information in the brain. Since the first recordings of electricity induced neurotransmitter release^1^ and of fast action potentials (APs) in peripheral nerves and neurons in the central nervous system^2,3^, the essential roles of APs in transducing signals between neurons and triggering neurotransmitter release have been well established. In addition, genetic, histological, and physiological studies have revealed each brain region to contain multiple neuronal classes with distinct electrical and chemical properties arranged in local and extended circuits^4^. How each node in a circuit relays or processes information will thus require the recording of APs in different cell types at high speeds and throughput, i.e. with low-millisecond temporal precision to enable accurate sequencing, and in enough neurons to acquire a representative information matrix. Furthermore, to understand single-cell operations such as integrating or filtering neurotransmitter inputs, large-scale recording of subthreshold potentials (SPs) will also be necessary.

To provide insight into these fundamental questions, the ability to record APs and SPs in large numbers of genetically defined neurons has long been desired. In the past two decades, extensive efforts have been made to improve the timing and sensitivity of AP detection by intracellular calcium reporters. APs open voltage-gated calcium channels to produce large, long-lasting increases in intracellular calcium, which can be reported by genetically encoded calcium indicators (GECIs) as increases in fluorescence. Notably, GECIs of the GCaMP family achieve high signal-to-noise ratios (SNRs) for spike bursts in vivo, and feature uniform responses across one- and two-photon excitation modalities. Conveniently, genetic encoding allows GECI expression to be restricted to particular cell subpopulations defined by lineage or connectivity, using cell type-specific promoters or viral injections to dendritic or axonal projection areas, or simply to be sparsified for discrete multi-unit recordings. GCaMPs are now widely used to determine whether a subpopulation of neurons is active during a sensation or behavior, where increased fluorescence indicates an increase in AP firing rates and thereby neuronal activation.

Unlike simply correlating neuronal activity to sensation or behavior, however, understanding calculations made by neuronal circuits requires kinetic precision. An example is the problem of how oscillations originate and entrain neurons. The gamma oscillation of 40 Hz arises during tasks that require attention or working memory, and its enhancement may improve long-term cognitive function ^5-7^. Measuring how spiking of specific subtypes of interneurons that generate the rhythm correlates with the firing of excitatory neurons will be necessary for understanding causal relationships between these rhythmic inputs and excitatory neuron threshold modulation ^8,9^. Timing is also important in the study of circuit plasticity, as backpropagating APs following synaptic inputs within 10 ms produce long-term potentiation (LTP), whereas backpropagating APs that precede synaptic inputs within 10 ms produce long-term depression (LTD)^10-12^. Thus, a change of a few milliseconds in AP timing can result in a change in plasticity direction, emphasizing the importance of timing. AP timings can be accurately assessed by extracellular electrode arrays, but these do not allow signal restriction to genetically specified cells.

Genetically encoded voltage indicators (GEVIs) provide an alternative method for AP recordings, combining the genetic targetability of GECIs with the temporal precision of electrodes. Voltage imaging has been performed at up to 3 kHz, with spike timing precision of <0.5 ms *in vivo*^13^. Meanwhile, the fastest GECI, jGCaMP8f, exhibits half-rise times to single APs of 6–7 ms *in vivo*^14^, and has been sampled at 200 Hz to achieve AP timing resolution of 7–10 ms. However, no published reports have compared GECI and GEVI responses in interneurons, whose frequent and brief APs pose a challenge to detection by either method.

Here we report the development and application of a high-performance GEVI, ASAP6c, for voltage imaging by one- or two-photon excitation in large numbers of densely labelled neurons in vivo. ASAP6c detects APs with 100% fluorescence increases and three-fold higher temporal precision than jGCaMP8f. In the mouse brain, ASAP6c detects APs in interneurons that are mostly missed by jGCaMP8f. These results with ASAP6c demonstrate the utility of high-performance voltage imaging for large-scale AP reporting with higher temporal resolution than possible with calcium indicators.

## Results

### Engineering positively tuned ASAPs for larger responsivity for action potentials

Previously, we developed the positively tuned ASAP4b and ASAP4e exhibiting high photostability^15^. While these had higher Δ*F*/*F*_0_ from –70 mV to +30 mV than the negatively tuned ASAP3 or ASAP5^13,16^, their onset kinetics are biexponential with only a minority component of the response exhibiting a time constant of < 10 ms. This could be due to steric or electrostatic interactions that cause resistance to upward S4 movement in ASAP4 variants. Thus, we set out to engineer positively tuned GEVIs with faster activation kinetics over ASAP4b and ASAP4e, while maintaining or improving steady-state responsivity.

As ASAP4 and ASAP3 differ primarily in linker sequences, we rescreened ASAP4.0, a precursor to ASAP4b and ASAP4e, for all amino acid substitutions at Gly-151 and at His-206 in two sequential rounds of electroporation-based screening (**Fig. 1a, Supplementary Note**). This identified ASAP4.0 G151K T206A as similar in responsivity to the previously reported ASAP4.0 T206H (ASAP4.1)^15^ (**Supplementary Fig. 1)**. In the third round, Phe-148 and Asn-149 double mutants were screened by electroporation and validated by patch-clamping (**Supplementary Fig. 2-3**). Two mutants, ASAP4.1 N149K and ASAP4.0 N149A G151K T206A, exhibited the largest SNRs to single APs **(Supplementary Table 1)**. In each round, we performed patch-clamp verification on the top 5-10 variants with better response and at least comparable kinetics to the parent template.

**Figure 1.**
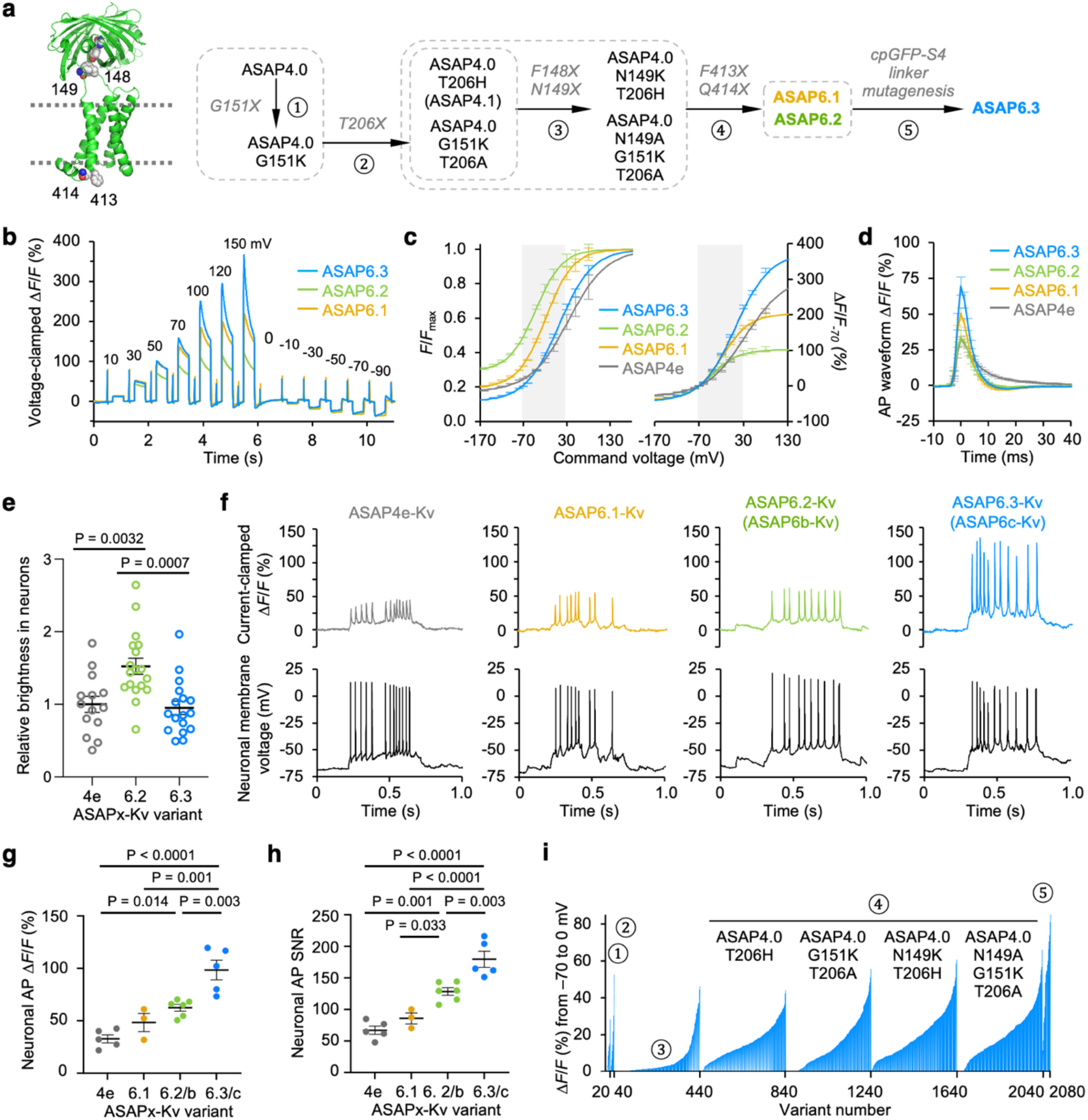
Protein engineering and *in vitro* characterization of the ASAP6 family of voltage indicators. **(a)** Left, ASAP4.1 protein model showing mutated positions. The engineered residues and the chromophore are highlighted as spheres. Right, engineering paths for ASAP6.1, ASAP6.2 and ASAP6.3. **(b)** Left, representative steady-state responses to voltage commands in HEK293A cells for ASAP6.1 - ASAP6.3 normalized to their fluorescence signals at -70mV. **(c)** F–V curves normalized to the maximum Δ*F*/*F* signal for each sensor (left) or to their signals at -70mV (right). N = 6, 4, 2, 5 for ASAP6.3, ASAP6.2, ASAP6.1, and ASAP4e. **(d)** Mean fluorescence responses to simulated AP-shaped waveform with 2ms half-width in HEK293A cells. N = 7, 4, 4, and 5 for ASAP6.3, ASAP6.2, ASAP6.1, and ASAP4e. **(e)** Relative brightness in rat cortical neurons for ASAP4e-Kv, ASAP6.2-Kv and ASAP6.3-Kv normalized to the mean value of ASAP4e-Kv. N = 14, 18, and 17 for ASAP4e-Kv, ASAP6.2-Kv, and ASAP6.3-Kv, respectively. **(f)** Representative optical traces (top) to evoked AP responses (bottom) recorded in cultured rat hippocampal neurons at physiological temperature at 800 fps. **(g, h)** Mean Δ*F*/*F* (g) and SNR (h) of optical APs. N = 5, 3, 7, and 6 neurons for ASAP4e-Kv, ASAP6.1-Kv, ASAP6.2-Kv, and ASAP6.3-Kv. **(i)**. Mean Δ*F*/*F* of all variants in the primary screen. Rounds correspond with numbering in **(a)**. Error bars represent standard error of the mean (SEM). Statistical differences were determined by one-way ANOVA with Tukey’s post hoc test.

A fourth round of mutagenesis focused on the voltage-sensing domain (VSD). In ASAP2s, a R414Q mutation on helix S4 increases steady-state responsivity but slows response kinetics^17^. R414Q removes a gating charge, which may reduce the electrostatic force on S4 upon depolarization, rendering mechanical barriers to S4 movement more difficult to overcome^18^. In addition, the amino acid 413 at the interface between the lipid bilayer and the cytosol tunes the F-V relationship in ASAP4.3–4.5. We thus generated and screened all 400 combinations of mutations at positions 413 and 414 in ASAP4.1 N149K and ASAP4.0 N149A G151K T206A (**Supplementary Fig. 4**). By increasing the imaging frame rate of our automated electroporation screening system to 300 frames-per-second (fps), we could obtain useful information on kinetics as well as steady-state responses from the same image series in the primary screen. To avoid being trapped in a local performance minimum specific to the third-round N149K mutation, we also performed F413X Q414X saturation mutagenesis on the second-round hit ASAP4.0 G151K T206A and the spectrally similar ASAP4.1 (**Supplementary Fig. 5**).

The four fourth-round screens revealed two interesting mutants (**Fig. 1b**). One, ASAP6.1, brightened 50% in response to 100-mV 2-ms-wide AP waveforms **(Supplementary Table 2**). Another, ASAP6.2, produced smaller responses of 34% per AP (**Supplementary Table 2)**. ASAP6.1 and ASAP6.2 exhibited mono-exponential time constants of 4.0 and 3.7 ms, respectively, at room-temperature (**Supplementary Table 3**). A single ≤4-ms time constant now accounted for 100% of the onset kinetics, as opposed to only 14% for the similar tau1 in ASAP4e.

To further improve responses, we performed a fifth engineering round, screening substitutions in the cpGFP-S4 linker of ASAP6.1 and ASAP6.2. One variant showed larger voltage responses than ASAP6.1 or ASAP6.2 (**Fig. 1b**) and was designated ASAP6.3. ASAP6.3 maintained the fast mono-exponential onset kinetics of ASAP6.2 (**Supplementary Table 3**) and exhibited the highest contrast of any GEVI to date, with a 7.9-fold response across all voltages and a fluorescence change (Δ*F*/*F*_0_) of 222% from –70 to +30 mV (**Fig. 1c**). ASAP6.3 also showed the largest Δ*F*/*F*_0_ to commanded AP waveforms among all variants (**Fig. 1d**), resulting in the highest predicted per-molecule SNR (**Supplementary Table 2)**.

### Comparison of ASAP4e-Kv, ASAP6b-Kv, and ASAP6c-Kv in neurons *in vitro* and *in vivo*

We assessed if ASAP6.3 brightness in neurons reached expected levels compared to earlier ASAP-family GEVIs. GEVIs with the same chromophore and chromophore pocket are expected to exhibit the same maximal molar fluorescence (m*F*_max_). If they are expressed to similar numbers per cell and the resting membrane potential in the cells is –70 mV, baseline cellular fluorescence should then track with fractional molar fluorescence at –70 mV (*mF*_–70mV_/*mF*_max_). In cortical neurons expressing GEVIs fused to the proximal retention and clustering sequence of Kv2.1 for somatic enrichment^19^, total cellular baseline brightness of ASAP6.2-Kv was similar to ASAP5-Kv, while ASAP4e and ASAP6.3 produced roughly half the brightness **(Fig 1e)**. This correlated with the *mF*_–70mV_/*mF*_max_ for each GEVI, revealing that ASAP6.3 was as well expressed in neurons as the other GEVIs.

We next tested the performance of the new ASAP variants in neurons. We expressed ASAP4e-Kv, -6.1-Kv, -6.2-Kv, and -6.3-Kv in primary rat hippocampal neurons and recorded fluorescence at 34-37 °C while evoking APs by plateau current injections. All variants faithfully recapitulated spike dynamics with sufficient SNR for spike trains as fast as 35 Hz in frequency (**Fig. 1f)**. Responses to single APs, measured as fractional change over baseline (Δ*F*/*F*_0_) averaged 98% for ASAP6.3-Kv compared to 62% for ASAP6.2-Kv (**Fig. 1g**), and SNR was also larger for ASAP6.3-Kv, reaching 2.7-fold of ASAP4e (**Fig. 1g, h**). ASAP6.3-Kv is also likely to report single APs with better SNR than ASAP5-Kv, as it exhibits 3-fold larger Δ*F*/*F*_0_ while maintaining 63% of baseline brightness (**Table S2**). On the other hand, ASAP6.1-Kv was inferior to ASAP6.2-Kv in Δ*F*/*F*_0_ and SNR in neurons (**Fig. 1g, h**), in contrast to its superiority in non-neuronal cells. Given these findings, we designated ASAP6.2 as ASAP6b for its high brightness, and ASAP6.3 as ASAP6c for its high contrast (Δ*F*/*F*_0_). The discovery of ASAP6b and ASAP6c involved the generation and screening of >2000 variants (**Fig. 1i**).

### Comparable performance of ASAP6c under one- and two-photon excitation

Previously characterized ASAP-family GEVIs respond similarly under one- and two-photon excitation^20^, but a comparison of responses under these two excitation regimes in the same set of labelled neurons in vivo has not previously been performed. We first characterized one-photon and two-photon excitation spectra of ASAP6c-Kv (**Fig. 2a**). We then expressed ASAP6c-Kv in pyramidal neurons of the CA1 hippocampus and imaged through a deep cranial window while the mouse was placed on a running wheel (**Fig. 2b**). Under widefield one-photon illumination at 800 fps, ASAP6c-Kv exhibited optical spikes with *ΔF*/*F*_0_ up to 100 %, consistent with the responses to APs observed in cultured neurons in vitro (**Fig. 2c**). Under resonant galvanometric two-photon scanning microscope at 500 fps, we also observed optical spikes with *ΔF*/*F*_0_ up to 100 % (**Fig. 2d**). These results confirmed that ASAP6c-Kv responds to spiking activity with similar amplitude during one- and two-photon imaging in vivo. Free-space Angular-Chirp-Enhanced Delay 2.0 (FACED 2.0) is an ultra-fast raster scanning two-photon imaging technique that enables kilohertz-rate imaging of hundreds of individual neurons in large 2D FOVs. FACED has been demonstrated before with the negatively tuned ASAP-family indicators ASAP3 at 920 nm and JEDI-2P at 1035 nm^21,22^, but has not yet been performed with a positively tuned voltage indicator. Both ASAP6c and JEDI-2P contain a T203H red-shifting mutation from Clover GFP^23^, and are similar in exhibiting peak excitation at 500-nm vs. the 485-nm peak of ASAP3 **(Fig. 2a)**. This shift allowed JEDI-2P to be excited with 1035-nm ytterbium fiber lasers, which produce pulses of high uniformity and power. We investigated if ASAP6c-Kv could also be efficiently imaged by FACED 2.0 using a 1035-nm laser. Indeed, ASAP6c-Kv revealed spikes in 385-Hz FACED 2.0 recordings with *ΔF*/*F*_0_ of ∼100% **(Fig. 2e)**. The SNR of these optical spikes, approximately 5, was similar to published values for JEDI-2P at similar illumination wavelengths and powers^22^. Thus, ASAP6c-Kv performs well with rapid two-photon scanning methods, including those using 1035-nm ytterbium fiber lasers.

**Figure 2.**
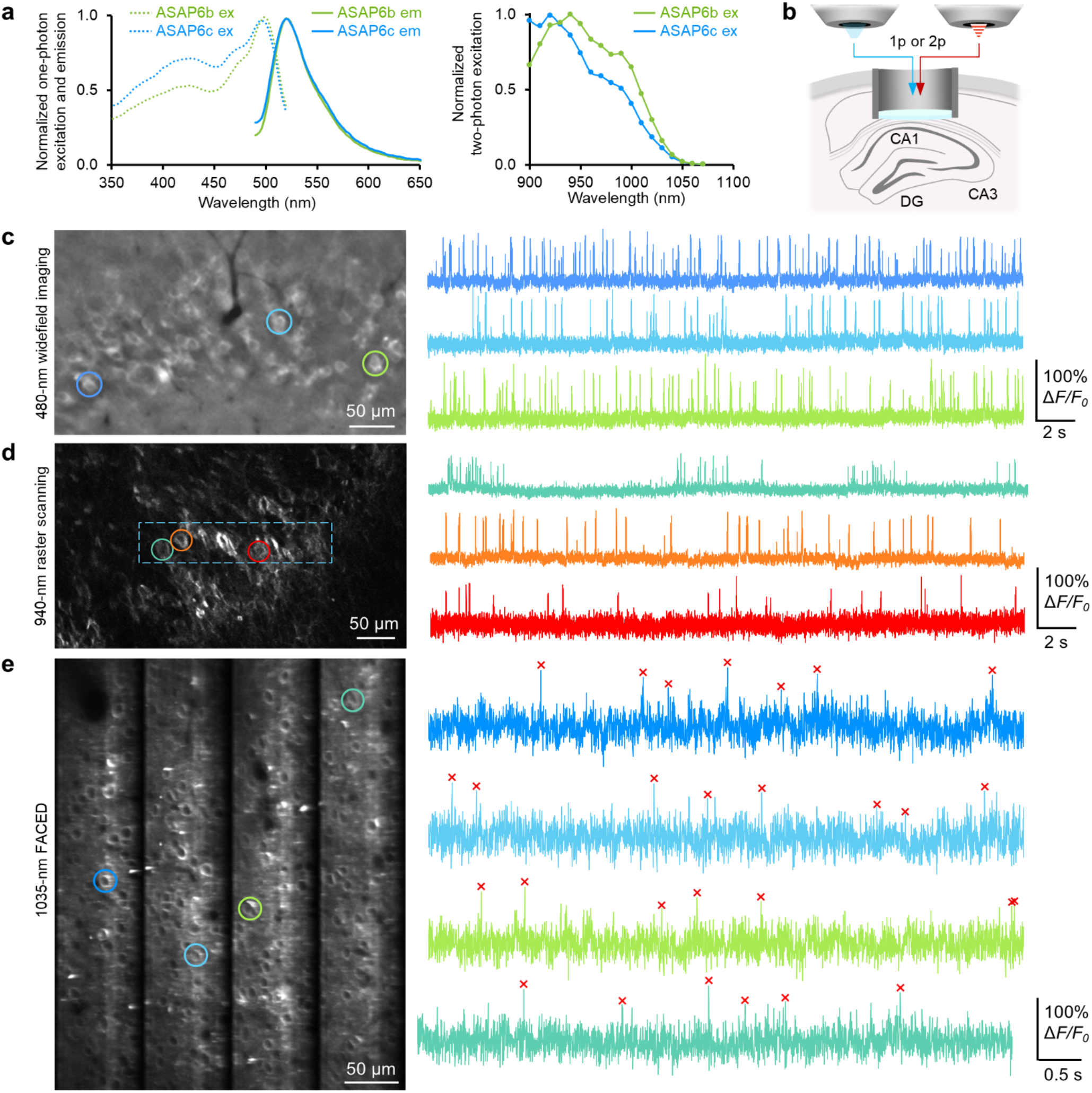
Similar responses by one and two-photon imaging in vivo, and fast 2p scanning. **(a)** Left, normalized one-photon excitation and emission spectra of ASAP6b and ASAP6c measured in HEK293-Kir2.1 cells. Right, normalized two-photon excitation measured in HEK293-Kir2.1 cells. **(b)** Schematic diagram of in vivo imaging in the CA1 hippocampus by one-photon or two-photon imaging. **(c)** One-photon voltage imaging of ASAP6c-Kv expressing CA1 pyramidal neurons from an awake-behaving mouse. The frames were acquired at 800 fps during widefield illumination at irradiance of 26 mW/mm^2^. The image on the left was frame-averaged after background subtraction and ΔF/F traces from neurons indicated with circles are shown on the right. **(d)** The same mouse from (c) was imaged using a two-photon microscope raster scanning with 940 nm light at 37 mW post-objective power. The image on the left was frame-averaged from a z-stack imaging result. The FOV indicated with a blue dotted rectangle was imaged at 500 fps for voltage imaging and the resulting ΔF/F traces from neurons indicated with circles are shown on the right. **(e)** Large-scale two-photon voltage imaging by FACED 2.0 system at 385 fps in the primary visual cortex of an awake-behaving mouse. ASAP6c-Kv expression was targeted to CaMK2α+ neurons in all cases.

### Superior performance of ASAP6c to earlier positively tuned GEVIs under one- and two-photon excitation

We compared ASAP4e-Kv, ASAP6b-Kv, ASAP6c-Kv, and pAce-Kv performance in mice by both one- and two-photon microscopy. In layer 2/3 primary motor cortex pyramidal neurons, all four GEVIs successfully showed spiking activity in awake-behaving mice when imaged at 400 fps (**Fig. 3a**). ASAP6b-Kv and ASAP6c-Kv optical spikes had higher SNR than those of ASAP4e-Kv and pAce-Kv **(Fig. 3b)**. In two-photon imaging with a conventional resonant galvanometric scanning microscope, all variants detected spikes in barrel cortex layer-2/3 neurons (**Fig. 3c)**. *ΔF*/*F*_0_ per AP was ∼80% for ASAP6c-Kv, ∼60% for ASAP6b-Kv, and ∼40% for ASAP4e-Kv, and SNR followed the same rank order **(Fig. 3d)**. Thus, ASAP6c-Kv demonstrates superior performance over ASAP6b-Kv, ASAP4e-Kv, and pAce-Kv under both one-photon and two-photon excitation in vivo, correlating with the characteristics of these GEVIs in vitro.

**Figure 3.**
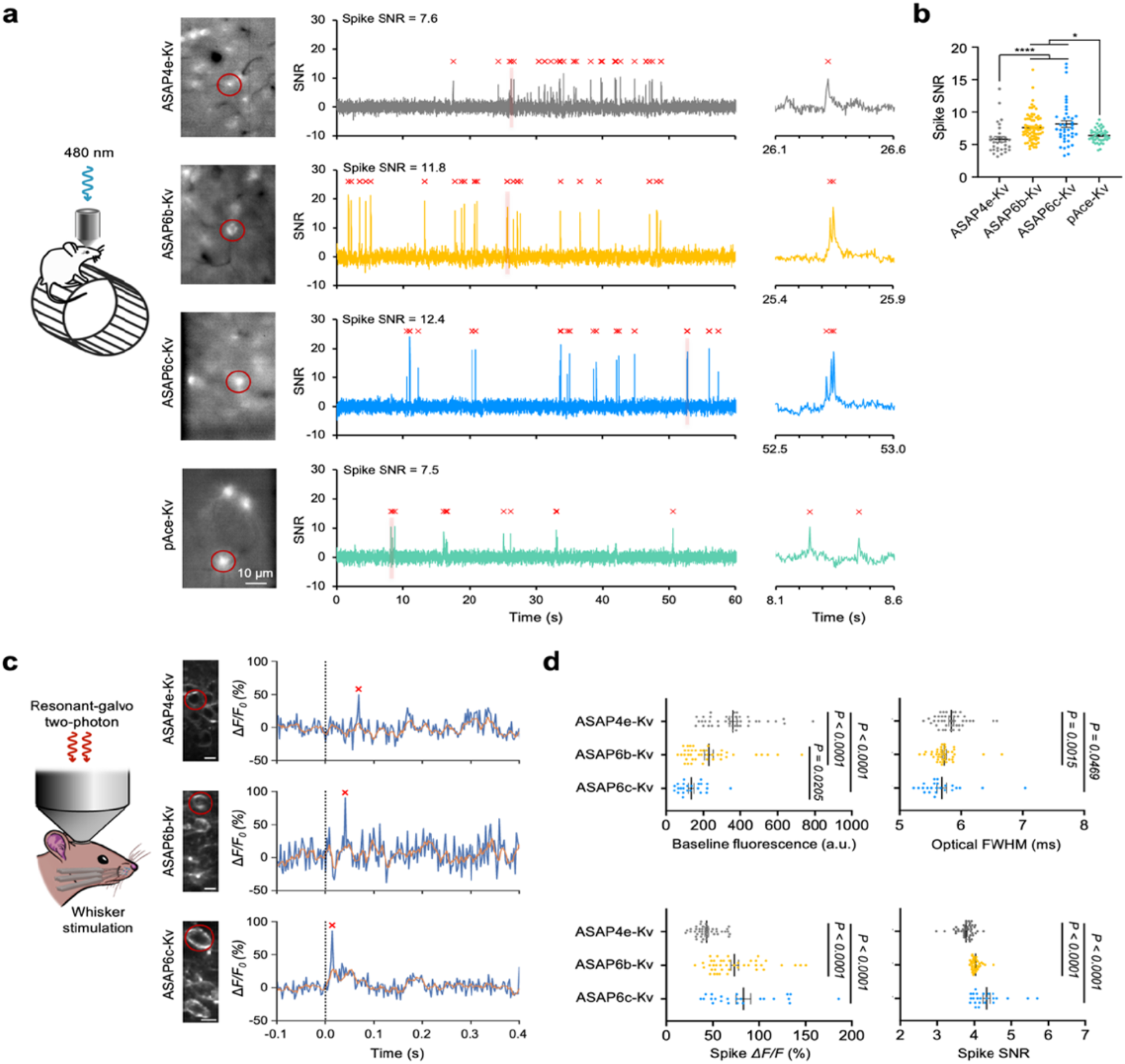
In vivo voltage imaging performance comparison by one- and two-photon microscopy. **(a)** One-photon comparison of ASAP4e-Kv, ASAP6b-Kv, ASAP6c-Kv, and pAce-Kv expressed in layer-2/3 pyramidal neurons in the mouse motor cortex. Left, One-photon fluorescence images of each sensor. The images were frame-averaged over 1-min. Middle, Example fluorescence traces from the neurons highlighted on the left with red circles. Red marks indicate optical spikes detected by a spike-detection algorithm, VolPy. The frame rate was 400 fps and imaged for 4 min at irradiance of 12 mW/mm^2^. The first 1 min segment is shown for display. Right, zoomed-in traces of the time window shaded in red on the traces in the middle. **(b)** Comparison of spike SNR. N = 32, 71, 40, and 43 neurons for ASAP4e-Kv, ASAP6b-Kv, ASAP6c-Kv, and pAce-Kv, respectively. Statistical differences were determined by Kruskal-Wallis test. The asterisks indicate p-values of <0.05 for *, and <0.0001 for ****. **(c)** Two-photon comparison of ASAP4e-Kv, ASAP6b-Kv, and ASAP6c-Kv expressed in mouse barrel cortex during whisker stimulation. 960 nm light was used for two-photon excitation and produced 40 mW post-objective power. Left, a schematic diagram. Middle, representative FOV for each sensor. Right, resulting fluorescence traces recorded at 400 fps. The dotted lines represent the timing of whisker-stimulation. The asterisks in red indicate detected spikes. Scale bars are 10 µm. (**d**) Statistical comparison of the three sensors for their baseline fluorescence, optical FWHM, spike *ΔF/F*_*0*_, and spike SNR. N = 42, 42, and 24 neurons were analyzed for ASAP4e-Kv, ASAP6b-Kv, and ASAP6c-Kv, respectively. Statistical differences were determined by Kruskal-Wallis test. P-values are shown above each comparison.

### Comparison of jGCaMP8f and ASAP6c-Kv for AP detection in interneurons in vivo

We next compared the ability of jGCaMP8f and ASAP6c-Kv to report APs in an interneuron subclass *in vivo*. APs in many interneuron types are challenging for both GECIs or GEVIs to detect due to their short durations and high frequencies. GEVIs require millisecond-scale activation kinetics to respond before the AP has passed, and both GECI and GEVI deactivation kinetics need to be fast enough so that closely spaced APs produce distinct and detectable peaks. We expressed jGCaMP8f or ASAP6c-Kv in interneurons marked by expression of Htr3a(BAC)-cre, which exhibit APs of 1.1–1.4 ms and firing frequencies up to ∼80 Hz^24,25^. We then performed one-photon imaging of labelled neurons in layer 2 (150–200 μm below the pia) of head-fixed awake mice at 500 fps and an equivalent irradiance of 16 mW/mm^2^ **(Fig. 4a)**. For both calcium and voltage movies, we used EXTRACT^26^ for signal source separation (**Fig. 4b,c**) followed by MLspike for AP detection. We chose MLspike as it has been used for both GECI-based and GEVI-based spike detection^13,27^, and performs well in this task even when compared to later algorithms^28^.

**Figure 4.**
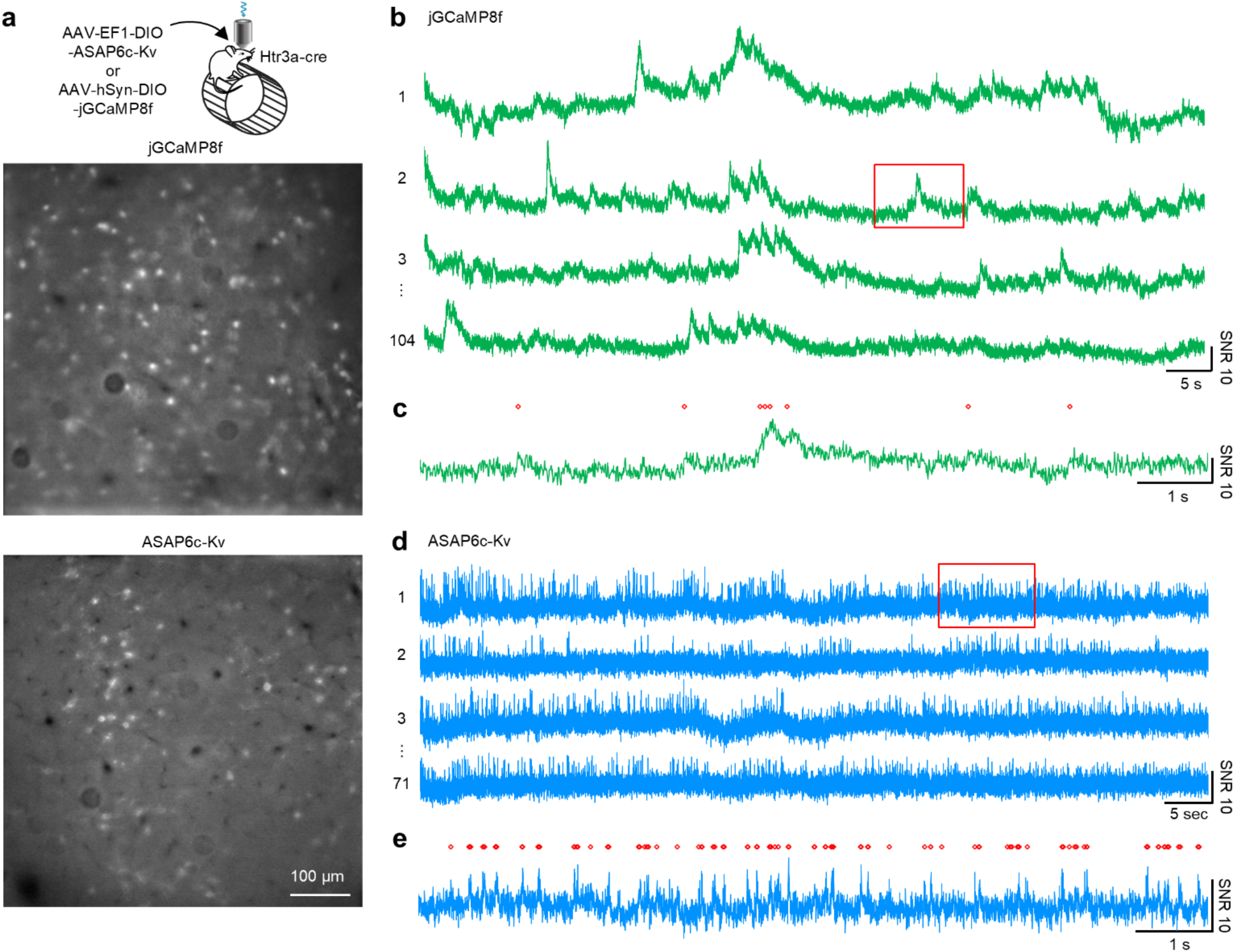
Improved spike detection by voltage imaging with ASAP6c-Kv compared to jGCaMP8f in Htr3a^+^ interneurons in vivo. **(a)** Top, a schematic diagram of one-photon in vivo imaging. AAVs of either EF1a-DIO-ASAP6c-Kv or hSyn-DIO-GCaMP8f were injected into the primary motor cortex of Htr3a^+^ cre driver mice. Bottom, frame-averaged images of each sensor expressed in Htr3a^+^ interneurons in L2/3 of M1. **(b)** Fluorescence traces of four jGCaMP8f expressing neurons. The image was acquired at 500 fps initially, but timebinned to 167 Hz for analyses. EXTRACT detected 104 ROIs in total. **(c)** Spikes detected by MLspike are indicated with red markers for the time period highlighted in red in (b). **(d)** Equivalent of (b) but shows ASAP6c-Kv traces at 500 Hz. EXTRACT detected 71 ROIs in total. **(e)** Equivalent of (c) but shows MLspike detected spikes for ASAP6c-Kv. All vertical scale bars show 10 SNR. Irradiance of 16mW/mm^2^ and 34 mW/mm^2^ were illuminated for GCaMP8f and ASAP6c-Kv expressing mice, respectively, but corresponding amplitude of baseline noise was added to ASAP6c-Kv traces to simulate effective irradiance of 16 mW/mm^2^ for both indicators.

Our results revealed multiple differences between jGCaMP8f and ASAP6c-Kv signals in Htr3a(BAC)-cre+ interneurons. EXTRACT identified similar numbers of neurons for the two indicators (50–100 per 1-mm^2^ field of view), but response amplitudes and event frequencies were strikingly different. jGCaMP8f responses ranged widely from 2 to 30 SNR units in amplitude, similar to published results in interneurons^14^. These variable responses can be explained by intracellular calcium reaching different levels depending on recent AP firing patterns^14^. In contrast, ASAP6c-Kv responses exhibited a tight range from 4 to 8 SNR units, as expected from its ability to closely track the voltage of each AP with <2-ms delays (**Fig. 4d,e**). Thus, in an interneuron subclass exhibiting high-frequency AP firing in vivo, ASAP6c-Kv was able to detect individual APs in a binary fashion with high SNR, whereas jGCaMP8f appeared to miss the majority of APs. Most APs modelled by MLspike from the jGCaMP8f trace were within long-lasting large-amplitude rises rather than associated with brief transients, suggesting difficulty in detecting isolated APs in this cell type using jGCaMP8f. These results are consistent with multiple published reports of absent responses of jGCaMP8f, jGCaMP8s, GCaMP6f, GCaMP6s, and XCaMP-G to most single APs in interneurons with ground truth provided by cell-attached patch recordings^14,28,29^. Thus, our results indicate that ASAP6c-Kv can report individual interneuron APs that escape detection by GECIs, including the recently developed fast GECI jGCaMP8f.

## Discussion

We introduce a positively tuned GEVI, ASAP6c, with higher Δ*F*/*F*_0_ and SNR for single APs than previous indicators^15^. Specifically, ASAP6c-Kv reports single APs with 100% Δ*F*/*F*_0_ in both one-photon and two-photon imaging in vivo. ASAP6c-Kv optical responses exhibit a narrow range of amplitudes even within bursts of APs, and can be resolved when separated by as little as 10 ms. Essentially ASAP6c-Kv detects APs in a binary fashion, similar to the all-or-none nature of APs themselves. In contrast, AP-induced calcium transients exhibit decay rates of >100 ms, which prevents accurate spike counting within closely spaced AP bursts. Indeed, we find that ASAP6c reports the high-frequency AP firing of interneurons whose individual APs are poorly detected by calcium imaging.

Opsin-based GEVIs can also detect electrical activity under two-photon illumination in some cases, but their responses exhibit complex dependencies on the illumination method and, so far, are smaller in SNR than ASAP-family indicators. Opsin-based GEVIs sense voltage only after light absorption drives the retinal-protein adduct into a transient voltage-sensitive photocycle intermediate^20^. The two-photon wavelength requirements for entering this voltage-sensitive state appears to differ between opsin-based GEVIs, with Voltron and Voltron2 requiring >1100 nm^30^, while pAce and Jarvis are voltage-responsive with 940 or 1030 nm excitation^31^. The half-life of this intermediate varies even between closely related opsins, with measurements of 46 ms for Voltron and 16 ms for Voltron2^30^, while a 2-fold drop in Δ*F*/*F*_0_ when excitation intervals are changed from 0.5 to 2 ms suggests a half-life of <2 ms for Jarvis^31^. In addition, Δ*F*/*F*_0_ of opsins are dependent on two-photon power density, and the requirement for high irradiance makes opsins less compatible with large-scale two-photon imaging using FACED or multi-target spiral scanning. For Voltron and Voltron2, Δ*F*/*F*_0_ increased with higher power densities (excitation 1135 nm, repetition rate 80 MHz, raster scanning), reaching 6% per AP at ∼40 mW post-objective power per layer-1 neuron^30^. Such powers limit the achievable number of targets^32^. In contrast to opsin-based GEVIs, Δ*F*/*F*_0_ of ASAP-family GEVIs is independent of irradiance in either one- or two-photon excitation. In addition, even an opsin-based GEVI optimized for two-photon responsiveness showed 33% lower SNR than the ASAP-family GEVI JEDI-2P when excited simultaneously in the same neurons in vivo^13^. Taken together, these observations reveal continuing limitations of opsin-based GEVIs under two-photon scanning illumination in response consistency and SNR.

Indeed, ASAP-family GEVIs, like GCaMPs, bring versatility to experimental planning as they respond well across various one- and two-photon imaging methods^31^. Voltage imaging in vivo requires achieving GEVI expression in a particular cell type and region by AAV packaging and empirical optimization of injection conditions or by generating transgenic reporter mice and crossing to driver mice. If different GEVIs are required for one- and two-photon imaging, then changing from one modality to the other would require repeating these steps with a different GEVI. The improved characteristics of ASAP6c-Kv in both one- and two-photon imaging allow laboratories performing voltage imaging to switch modalities seamlessly, and even use the same animal preparation to test both one- and two-photon imaging methods, as performed here (**Fig. 2**), to determine which is more suitable for the experimental question being investigated.

In summary, ASAP6c-Kv offers neuroscientists the ability to record spiking activity in genetically defined neuronal populations in vivo at high rates, high density, and various depths with one- or two-photon microscopy, enabling studies of how fast electrical signaling between neurons perform calculations and convey information in the brain.

## METHODS

### Sensor engineering and *in vitro* characterization

#### Plasmid construction

For electroporation-based screening transfection into HEK293-Kir2.1 Cells for electrical screening, ASAP variants were subcloned into pcDNA3.1 with a CMV enhancer and promoter and bGH poly(A) signal. All plasmids were made by standard molecular biology techniques followed by a Sanger sequencing (Sequetech). PCR fragment libraries were generated using standard PCR techniques, which were used to directly transfect HEK293-Kir2.1 cells with the linear product using lipofectamine 3000 (Thermo Fisher Scientific). For characterization in HEK293A cells, all voltage indicator variants were subcloned into a pcDNA3.1/Puro-CAG vector ^23^. For *in vitro* characterization in cultured neurons, ASAP variants were subcloned into pAAV.hSyn.WPRE vector. The soma-targeting tag from Kv2.1 was used as previously described to restrict ASAP expression in soma and proximal dendrites^19,33^.

#### Cell lines

All cell lines were maintained in a humidified incubator at 37°C with 5% CO_2_. For electroporation-based screening, HEK293-Kir2.1 cell line was maintained in high-glucose DMEM, 5% FBS, 2 mM L-glutamine, and 500 μg/mL Geneticin (Life Technologies) as previously described^13^. For patch-clamp recordings for *in vitro* characterization, HEK293A cells were used and cultured in the same DMEM based media as described above but without adding Geneticin.

#### Primary neuronal culture and transfection

Hippocampal neurons were isolated from the Hippocampi of embryonic day 18 Sprague Dawley rats by dissociating in TrypLE™ Select Enzyme (Gibco) supplemented with 0.005% DNaseI in an incubator maintained at 37°C and 5% CO_2_. The dissociated neurons were plated on poly-D-lysine coated 12 mm diameter glass coverslips at a density of 4 × 10^4^ cells/cm_2_. Cells were plated for several hours in Neurobasal media (Gibco) with 10% v/v FBS (Gemini bio), 2mM GlutaMAX (Gibco), and 5% v/v B27 supplement (Gibco), then the media were replaced with Neurobasal media supplemented with 1% FBS, 2mM GlutaMAX, and B27. A half of the media in each well was replaced every 3 days with fresh Neurobasal media supplemented with the same GlutaMAX and B-27, but without FBS. 5-Fluoro-2’-deoxyuridine (Sigma-Aldrich) was typically added at a final concentration of 16 uM at DIV 4–7 to limit non-neuronal cell growth. On DIV 7–10, Hippocampal neurons were transfected using a modified Lipofectamine 2000 (Life Technologies) transfection procedure as previously described^15^.

#### Viruses

All ASAP viruses were produced by the Stanford Neuroscience Gene Vector and Virus Core facility at Stanford University. AAV9-syn-FLEX-jGCaMP8f-WPRE and AAV9.CamKII0.4.Cre.SV40 were purchased from Addgene.

#### High-throughput automated electroporation-based cell screening

Electroporation-based cell screening was conducted following the procedure as previously described^15^. Briefly, HEK293-Kir2.1 cells were plated in 384-well bottomless adhesive plates (Grace Bio-Labs) attached with an indium tin oxide coated conductive glass slides (Delta Technologies). Poly-D-lysine hydrobromide (MP biomedicals) was coated before plating the cells. ASAP mutation libraries were PCR amplicons with a CMV enhancer and promoter at 5’ end and bGH poly(A) signal at 3’ end as previously described^34^. Cells were transfected with either plasmid controls or PCR fragment libraries in 384-well plates with Lipofectamine 3000 (∼100 ng plasmid DNA or equivalent transfection efficiency of PCR fragments, 0.4 μL p3000 reagent, 0.4 μL Lipofectamine per well) followed by a media change 4–5 hours later. The screening was done 2 days post-transfection in Hank’s Balanced Salt solution (HBSS) buffered with 10 mM HEPES (Life Technologies). Cells were imaged at room temperature on an IX-81 inverted microscope using a 20x 0.75-numerical aperture (NA) objective lens (all Olympus). A 120-W metal halide lamp (X-Cite 120PC, Exfo) served as the excitation light source. The filter cube set consisted of a 480/40nm excitation filter and a 503-nm long pass emission filter. During the screening, an operator located the best field-of-view (FOV) and the focus for each well. A single FOV per well was imaged for a total of 3 s, with a 10-μs 150-V square pulse applied near the 1-s mark. The fluorescence was collected at 300 Hz (3.3-ms exposure time per frame) by an ORCA Flash4.0 V2 C11440–22CA sCMOS camera (Hamamatsu) with the FOV cropped to 512×170 pixels (binned 4×4). Each mutant had at least 4 replicates.

#### Whole-cell patch-clamping and imaging of HEK293a cells

Patch-clamp experiments were conducted as previously described^15^. Briefly, the transfected HEK293A cells were replated on 12-mm diameter glass coverslips (Carolina Biological), and each coverslip was located in a patch chamber after 24 hours of transfection. Following a successful giga-ohm seal with a patch pipette with resistance of 3–5 MΩ, the cell was patched in a whole-cell voltage-clamp mode with a Multiclamp 700B amplifier controlled by pClamp software (Molecular Devices). Fluorophores were illuminated with UHP-F-455 LED (Prizmatix) passed through a 484/15-nm excitation filter (Chroma) and focused on the sample through a 40× 1.3-NA oil-immersion objective (Zeiss). Emitted fluorescence passed through a 525/50-nm emission filter (Semrock) and was captured by an iXon 860 electron-multiplying charge-coupled device camera (Oxford Instruments) cooled to maintain –80°C. For all experiments fluorescence traces were corrected for photobleaching by dividing the recorded signal by an exponential fit to the data. To characterize steady-state fluorescence responses, cells were voltage-clamped at a holding potential of –70 mV, then voltage steps of 1-s duration between –120 and +120 mV were imposed. To obtain complete voltage-tuning curves, voltage steps from –200 to +120 mV were used. When the membrane potential could not be readily clamped for some of the less physiological voltage steps, the relevant points were excluded from data analysis. A scaled action potential waveform (FWHM = 3.5 ms, –70 to +30mV) recorded from cultured hippocampal neuron was included before each step to estimate the fluorescence change of the indicators to action potential waveform.

The kinetics of ASAP6 variants were calculated from voltage imaging data recorded at 800Hz camera frame rates in room-temperature. Fluorophores were illuminated with SOLIS-470c LED (Prizmatix) passed through a 490/20-nm excitation filter (Chroma) and focused on the sample through a 40× 1.3-NA oil-immersion objective (Zeiss). Emitted fluorescence passed through a 540/50-nm emission filter (Semrock). The ASAP expressing HEK293A cells were patched in a whole-cell voltage clamp mode and was given a pulse protocol to command the membrane potential change. The fluorescence response to a 100-mV step pulse was fitted to a mono-exponential function for both onset (depolarization) and offset (repolarization) segments to determine the relevant time constants. To test ASAP6 variants’ responses to simulated AP-shaped waveforms, a spontaneous spike with 100-mV amplitude recorded from a cultured hippocampal neuron was modified to have either 2ms or 4ms full-width at half-maximal (FWHM) and used as a command voltage for the HEK293A cells expressing ASAP6 variants. The fluorescence images were acquired at 800Hz.

#### Whole-cell patch-clamping and imaging of cultured hippocampal neurons

Cultured rat hippocampal neurons were transfected with 150 ng of corresponding ASAPx-Kv plasmids under the hSyn promoter on DIV 10 as described above. Patch-clamp electrophysiology was then performed 4–5 days post-transfection following the procedure described in ^16^. A coverslip with cultured neurons was kept in a patch chamber on the stage of Axiovert 100M inverted microscope (Zeiss), and were continuously perfused with extracellular solution prepared with 145 mM NaCl, 3 mM KCl, 2 mM CaCl_2_, 2 mM MgCl_2_, 10 mM HEPES, 10 mM glucose, and pH was adjusted to 7.4 with osmolarity of 310 mOsm/kg. The solution was heated to produce 34–37 °C in the patch chamber. Patch pipettes were filled with 123 mM K-gluconate, 10 mM KCl, 8 mM NaCl, 1 mM MgCl_2_, 10 mM HEPES, 1 mM EGTA, 0.1 mM CaCl_2_, 1.5 mM MgATP, 0.2 mM Na_4_GTP, 4 mM glucose, and pH was adjusted to 7.2 with osmolarity of 300 mOsm/kg. Action potentials were evoked by injecting step currents in whole cell current clamp mode and recorded at the sampling rate of 10 kHz. Starting from 0 pA, current steps were applied for 11 sweeps with 20-pA increments with the pulse interval of 0.5 sec. Neurons requiring more than ± 100 pA holding current to maintain resting membrane potential near –70 mV were discarded. For voltage imaging of cultured neurons, the blue light from SOLIS-470c LED (Thorlabs) was filtered through a 490/20-nm band-pass filter (Chroma) and focused by a 40× 1.3-NA oil-immersion objective lens (Zeiss). The power density at the sample was 55 mW/mm^2^. The resulting fluorescence emission filtered by a 540/50-nm bandpass filter (Semrock) was captured at 800 fps using iXon 860 EMCCD (Andor). Fluorescence traces were corrected for photobleaching and normalized to baseline by using custom codes in MATLAB as previously described^15^, then exported to Excel (Microsoft) for graphing.

#### One-photon spectra measurement in HEK293-Kir2.1 cells

HEK293-Kir2.1 cells were maintained and transfected as previously described in Hao and Lee et al. ^16^ with the following modifications. Two days before measurement, HEK293-Kir2.1 cells were plated in 24-well glass-bottom plates at 180,000 cells per well. Three wells were prepared for each sensor variant.

After 12 hours of incubation, cells in each well were transfected with the corresponding plasmid DNA in pCaggs backbone vector using Lipofectamine 3000. Specifically, 400 ng of plasmid DNA, 1 μL of P3000 reagent, and 1 μL of Lipofectamine 3000 were pre-mixed in 50 μL of Opti-MEM medium. The mixture was incubated at room temperature for 10 minutes, then added to each well. Following overnight incubation, the medium was replaced with fresh culture medium and cells were incubated until measurement.

Two days after transfection and immediately before spectral measurement, the culture medium was replaced with 100 μL of 10 mM HEPES-supplemented HBSS per well. Cells were detached from the glass surface and dissociated by gentle pipetting. Subsequently, the 100 μL cell suspension was transferred to a 96-well glass-bottom plate for measurements.

Spectral measurements were conducted using a Varioskan microplate reader (Thermo Fisher Scientific). Excitation spectra were recorded across the 300–520 nm range while monitoring emission at 545 nm. Emission spectra were recorded across the 490–700 nm range with excitation at 470 nm. Increments of 2 nm were used for both measurements. Results from three replicates were averaged for each sensor variant, then normalized to maximum signal before smoothing with a 5-point running average.

### Hippocampal voltage imaging

#### Virus injection and deep hippocampal window surgery

The intracranial virus injection in the Hippocampus CA1 was performed following the procedure described in ‘One-photon voltage imaging in motor cortex excitatory neurons’ section with following modifications. Female mice aged 12-13 weeks were injected with AAV9-EF1a-DIO-ASAP6c-Kv (2.2E+13 vg/mL) and AAV9.CamKII0.4.Cre.SV40 (2.3E+13 vg/mL). Each virus was pre-diluted, such that there was 3.5E+8 vg of the ASAP6c-Kv virus and 1.1E+7 vg of the Cre virus in the final injected volume (250 nL). The injection was done on the right hemisphere at AP : -2.0, ML : 1.5, and DV : 1.8 (all measured from bregma) to target the cell body layer of CA1. Two to four weeks after the injection, a deep cranial window was installed following the procedure described previously ^15^.

#### One-photon hippocampal imaging

For one-photon single plane voltage imaging in CA1, a BX-51 microscope (Olympus) equipped with a 505-nm LED (SOLIS-505c, Thorlabs), a long-working distance 16× objective lens (NA 0.8, Nikon), a 0.63× demagnifying c-mount adapter, a commercial sCMOS camera, 480/40nm bandpass filter (Chroma; excitation), and a 540/50nm bandpass filter (Semrock; emission) were used. Two weeks after the cranial window surgery, the mice were handled 5-10 minutes for two days. Then the mice were head-fixed and placed on a running wheel for five days by gradually increasing the duration from 10 to 30 minutes for habituation. For imaging, the light intensity was kept at 26 mW/mm^2^ during the 60-sec imaging sessions for each FOV. The fluorescence images were captured at 800 fps on an sCMOS camera. Resulting images were first corrected for lateral motions by using NormCorre package ^35^. Cellular ROIs were identified manually by using Fiji. No additional bleach correction or denoising was performed for the results shown in Figure 2c.

#### Two-photon hippocampal imaging

Two-photon in vivo voltage imaging in CA1 was performed using a custom-built resonant-Galvanometric two-photon setup as previously described ^16^. Briefly, the head-fixation and running wheel platform was moved to the two-photon microscope rig to keep the mice under the habituated environment during imaging. A mode-locked tunable ultrafast laser (InSightX3, Spectra-Physics) at 940 nm was used for imaging with a post-objective power of 37 mW. A 211×33-pixel field of view was imaged at 500 fps for 2 min. Resulting images were first corrected for lateral motions by using NormCorre package ^35^. Cellular ROIs were identified manually by using Fiji. After confirming that there was no bleed-through signal from each neuron, pixels adjacent to each cellular ROI were chosen to be used for subtracting the background signal. Then the mean fluorescence from the frames corresponding to 1-2 sec of each trace was used for normalization to acquire Δ*F/F*_0_ traces. No additional bleach correction or denoising was performed for the results shown in **Fig. 2d**.

### Two-photon *in vivo* imaging in visual cortex (FACED 2.0)

All animal experiments were carried out in accordance with the National Institutes of Health guidelines for animal research. The procedures and protocols involving mice were approved by the Institutional Animal Care and Use Committee at the University of California, Berkeley.

Virus injection and cranial window implantation were performed as previously described^36^. Briefly, wild-type C57BL/6J mice were anesthetized with isoflurane (1–2% in oxygen) and administered 0.3 mg/kg buprenorphine subcutaneously. Animals were head-fixed in a stereotaxic apparatus (Model 1900, David Kopf Instruments), and a 3.5-mm craniotomy was made over the left visual cortex while leaving the dura intact. A glass pipette (inner diameter 15–20 µm, 45° bevel) controlled by a hydraulic manipulator (MO10, Narishige) was used to inject a mixture of CaMKIIα-Cre and ASAP6c-Kv at 6–12 sites (20–50 nL per site) at a depth of 250 µm in the left visual cortex. A glass coverslip (No. 1.5, Fisher Scientific) was embedded into the craniotomy and sealed with dental acrylic. A titanium headpost was then attached to the skull using cyanoacrylate glue and dental acrylic. *In vivo* imaging was performed after 3 weeks of recovery in head-fixed, awake mice.

We tested ASAP6 for population voltage imaging using an ultrafast scanning two-photon fluorescence microscope based on free-space angular-chirp-enhanced delay (FACED), as previously described^21,22^. Briefly, the output of a 1035-nm fiber laser (Monaco 1035-40-40, 1-MHz repetition rate; Coherent) was focused by a cylindrical lens and directed onto a pair of mirrors. Through free-space angular-chirp-enhanced delay (FACED), each laser pulse was split into ∼100 spatially separated and temporally delayed pulses, generating at the focal plane of a 25×1.05 NA objective (XLPLN25XWMP2, Olympus) a line scan along X axis with a 0.8-μm pixel size and 1-ns inter-pulse delay at a rate of 1 MHz. A galvo scanning along the y axis at 769.2 Hz generated an 80×400-µm FACED FOV at a frame rate of 1.5 kHz. An additional x-axis galvo tiled multiple FACED FOVs to image a 320×400-µm FOV at 384.6 Hz or a 160×400-µm FOV at 769.2 Hz. The post-objective excitation power was 175 mW.

We recorded voltage activity from ASAP-expressing L2/3 neurons in the primary visual cortex of a head-fixed awake mouse during the presentation of drifting grating visual stimulation. Visual stimulation was programmed in MATLAB (MathWorks) using Psychophysics Toolbox^37^. Drifting gratings were presented on a monitor, with each stimulation cycle consisting of a 0.5-second blank period followed by a 0.5-second drifting grating. Grating orientations ranged from 0° to 315° in 45° increments. Each trial included all eight orientations and lasted 8 seconds. 10, 15, or 20 repeated trials were acquired for each FOV.

Data were analyzed using custom programs in MATLAB (Mathworks). Time streaming images were motion registered using an iterative cross-correlation method^1^. ROIs were selected using a combination of automatic 2D cell segmentation (Cellpose 2.0)^38^ and manual correction. Then, the average signal within each ROI was extracted as the raw neuronal trace. To calculate ΔF/F, the baseline signal (F) was obtained by averaging the signal from 0.3 to 0.5 seconds during the blank period, and ΔF was defined as the difference between the raw trace and F. For spike detection, the ΔF/F trace was detrended, and peaks exceeding 3.2 times the sliding standard deviation (std) of the detrended ΔF/F were identified as valid spikes.

### One-photon voltage imaging in motor cortex excitatory neurons

#### Surgery and viral injection

To prepare mice for one-photon *in vivo* imaging from primary motor cortex, we followed the procedure previously described^15,39^ with a few modifications. Briefly, 5–6 weeks old male wild-type C57BL/6J mice (Jackson Laboratories, No. 000664) were injected with relevant GEVI viruses (AAV9-EF1a-DIO-ASAP4e-Kv-WPRE (5.9E+12 vg/mL), AAV9-EF1a-DIO-ASAP6b-Kv-WPRE (4.0E+13 vg/mL), AAV9-EF1a-DIO-ASAP6c-Kv-WPRE (2.3E+13 vg/mL), or AAV9-EF1a-DIO-pAce-Kv-WPRE (2.1E+13 vg/mL)) mixed with CaMKIIα Cre virus (2.3×10^13^ GC/mL, Addgene #105558). For each mouse, 450 nL of the GEVI virus pre-diluted to match the lowest titer among the four GEVI viruses (that is, 5.9E+12 vg/mL) was mixed with 50 nL of CaMKIIα Cre virus pre-diluted to make the GEVI virus (floxed) to diluted CaMKIIα Cre virus ratio to be 40:1. The intracranial virus injection was done on the left hemisphere at the coordinate; ML –1.5 mm, AP 1.0 mm, and DV 1.2–1.0 mm (all measured from bregma), for labeling of layer 2/3 pyramidal neurons in motor cortex. Cranial window surgery was done 8–9 weeks after the injection. In brief, a 3×3-mm craniotomy was cut using #11 scalpel blades (Fine Science Tools) and a square coverslip (#1 coverslip glass, Warner Instruments) was implanted on top of the dura within the craniotomy with mild compression of the brain. The window was centered at ML –2.0 mm, AP 0.5 mm. The window was sealed to the skull using dental cement (C&B Metabond, Parkell) and at the end of surgery, a custom titanium headplate was attached to the skull to fixate the head to the microscope stage during subsequent imaging. Once the animals recovered, and the cranial window and head-plate stayed in good quality, all the mice were made blind to the experimenter by another researcher. The experimenter was left blind for the identity of each animal until all one-photon in vivo imaging and spike analyses were finished.

#### One-photon *in vivo* imaging and analysis

One-photon widefield voltage imaging of head-fixed mice was performed as previously described^15^. Briefly, a BX-51 microscope (Olympus) equipped with a 505-nm LED (SOLIS-505c, Thorlabs), a long-working distance 16× objective lens (NA 0.8, Nikon), a 0.63× demagnifying c-mount adapter, a Flash4.0 V2 scientific CMOS (sCMOS) camera (C11440-22CA, Hamamatsu), 480/40nm bandpass filter (Chroma; excitation), and a 540/50nm bandpass filter (Semrock; emission) were used. High-speed acquisition of fluorescence images at 400 fps was achieved by reading 128 vertical lines and 512 horizontal lines from the 4×4-binned sCMOS sensor. The light intensity was kept at 12 mW/mm^2^ during the four-minute imaging sessions for each FOV. Resulting images were corrected for lateral motions using NormCorre package^35^. The ROIs for neurons were selected using a graphical user interface (GUI) within VolPy package. Spike locations were determined using the spikepursuit algorithm in VolPy^40^. Using the peak signal amplitude of the located spikes, the SNR of each event was calculated in VolPy.

### Two-photon *in vivo* imaging in barrel cortex

#### Virus injection and cortical window surgery

Mice aged two to three months old were anesthetized with isoflurane (1.0–1.5% in O2). C1, C2 and C3 whisker columns were localized using transcranial intrinsic signal optical imaging to target the viral injections. A 3-mm diameter craniotomy was made centered on the C2 column^41^. AA9-hSyn-ASAP6-kv was injected at 5–6 locations surrounding the C2 column at 250 µm and 350 µm depth. A chronic cranial window (3-mm diameter glass coverslip, No. 1 thickness, CS-3R, Warner Instrument) was attached with dental cement after viral injections. Before surgery, mice received dexamethasone (2 mg/kg), enrofloxican (5 mg/kg), and meloxicam (10 mg/kg). Buprenorphine (0.1 mg/kg) was administered as the post-operative analgesic.

#### Stimulus Delivery

At the start of each imaging session, mice were placed under light anesthesia with isoflurane and head-fixed under the 2-photon microscope. 9 whiskers (rows B-D, arcs 1-3) were inserted into a 3×3 array of small tubes with calibrated piezoelectric actuators, centered on the C2 whisker. Trials consisted of single 1×250-μm or trains of 5×150-μm whisker deflections that were applied 5 mm from the face with 100-ms spacing between stimuli within a train. Whisker stimuli were delivered in random order with equal probability. An inter-trial interval of 1.5 ± 0.5 s was used between trials. Approximately 80 trials were delivered over the course of 15 minutes imaging. Paw guards prevented paw contact with the piezos. Imaging experiments were performed in total darkness. Uniform white noise was continuously applied to mask sounds from piezo actuators. Stimulus delivery was controlled by an Arduino Mega 2560 and custom routines in Igor Pro (WaveMetrics).

#### Two photon imaging

Two-photon imaging took place 3–4 weeks after viral injection on awake head-fixed mice. Imaging was performed with a modified, commercially available (ThorLabs) two-photon microscope^42^. A titanium-sapphire laser (Chameleon Ultra II, Coherent) was used as the two-photon excitation source, and a wavelength of 960 nm was used to excite ASAP6-Kv. A post-objective power of 30–50 mW was used for imaging at depths of 160–200 µm below brain surface. 192×64- or 96×64-pixel fields of view were imaged using a pixel size of 0.694 µm per pixel or 0.798 µm per pixel, resulting in a 133×34- or 77×52-μm area, respectively, imaged at 220 fps.

#### Image Processing

Fluorescence images were corrected for x-y motion using NoRMCorre^35^. Neuronal regions-of-interests (ROIs) were manually annotated using ImageJ (NIH) based on the mean intensity image. Δ*F*/*F*_0_ traces were extracted, with *F*_0_ defined as the mode of the fluorescence histogram of the ROI across each 15-minute movie. Spike locations and subthreshold activity was extracted using the CaImAn algorithm^40^ and custom Python routines (available at the <u>SpkXtract</u> Github repository) unless otherwise stated.

### One-photon voltage and calcium imaging in Htr3a+ interneurons in motor cortex

#### Virus injection and cortical window surgery

The virus injection and cranial window surgery were performed following the procedure described in ‘One-photon voltage imaging in motor cortex excitatory neurons’ section with following modifications. Seven to eight weeks old female TG(Htr3a-cre)NO152Gsat mice were injected with 500 nL of either AAV9-EF1a-DIO-ASAP6c-Kv (2.2E+13 vg/mL) or AAV9-syn-FLEX-jGCaMP8f-WPRE (3.0E+13 vg/mL) in the left hemisphere at ML : -1.5, AP : 1.0, and DV : 1.2–1.0 (all measured from bregma). Cranial window surgery was performed 3 weeks after injection.

#### One-photon cortical imaging and analysis

One-photon widefield voltage imaging of head-fixed mice was performed following the procedure described in ‘One-photon voltage imaging in motor cortex excitatory neurons’ section with following modifications. Briefly, a BX-51 microscope (Olympus) equipped with a 505-nm LED (SOLIS-505c, Thorlabs), a long-working distance 16× objective lens (NA 0.8, Nikon), a 0.63× demagnifying c-mount adapter, a commercial sCMOS camera, 480/40nm bandpass filter (Chroma; excitation), and a 540/50nm bandpass filter (Semrock; emission) were used. The light intensity was kept at either 16 mW/mm^2^ for jGCaMP8f or 34 mW/mm^2^ for ASAP6c-Kv during the 90-sec imaging sessions for each FOV. The fluorescence images were captured at 500 fps. Resulting images were first corrected for lateral motions by using NormCorre package ^35^. Then the ROIs and corresponding fluorescence traces were extracted by using the automated cell-extraction algorithm EXTRACT ^26^. To estimate the spike locations, we used MLspike for both ASAP6c-Kv and jGCaMP8f traces ^43^. We decreased the effective exposure of ASAP6c-Kv images (adding noise to simulate 2.2-fold lower exposures) to compensate for 2.2-fold higher excitation powers used for ASAP6c-Kv vs jGCaMP8f acquisition. Likewise, the jGCaMP8f signal was time-binned by a factor of 3 by averaging every 3 frame to simulate 167 fps from an original imaging result of 500 fps, improving SNR over faster framerates while capturing multiple timepoints on the upward portion of the jGCaMP8f response^14^.

## Declaration of interests

S.L., Y.A.H., T.R.C., and M.Z.L. are inventors on a patent for the earlier ASAP5 voltage indicator. The remaining authors declare no competing interests.

## Acknowledgements

This work was supported by National Institutes of Health (NIH) Grants 1R01MH126904-01A1 (L.M.G.), R01MH130452 (L.M.G.), U19NS118284 (L.M.G.), U01NS118300 (N.J.), U01NS137449 (N.J.), R21DC020325 (M.Z.L.), R01NS123681 (M.Z.L., J.D., L.M.G., T.R.C., N.J.), UM1MH136462 (M.Z.L.), and 1RM1NS132981 (M.Z.L.), and by a Wu Tsai Neurosciences Institute Big Ideas Grant (M.Z.L.). Additionally, N.J. acknowledges support from the Weill Neurohub, L.M.G. is an HHMI Investigator, and T.R.C. is a Chan-Zuckerberg Biohub Investigator. We thank U. Afifa (University of California, Berkeley) for experimental assistance; P. Rupprecht (University of Zurich), F. Dinç (Stanford University), and M. Schnitzer (Stanford University) for advice on signal analysis; and I. Soltesz (Stanford University) for feedback on the manuscript.

## Author contributions

S.L. engineered and characterized ASAP6c in vitro, performed in vitro brightness and photostability tests, prepared primary neuron cultures, conducted all patch clamp experiments in neurons, conducted mouse surgeries, imaging, and data analyses for one-photon in vivo imaging of pyramidal neurons in motor cortex, one-photon in vivo imaging of interneurons in motor cortex, and one- and two-photon in vivo imaging of CA1 pyramidal neurons in Hippocampus, prepared figures, and co-wrote the manuscript. G.Z. engineered and characterized ASAP6.1 and ASAP6b in vitro, prepared figures, and co-wrote the manuscript. Y.A.H. performed data processing and analysis of the multi-plane one-photon imaging of CA1 pyramidal neurons, wrote codes for data analyses, prepared figures, and co-wrote the paper. R.H.R. conducted mouse surgeries for one-photon in vivo imaging of interneurons in motor cortex, provided guidance for one- and two-photon in vivo imaging performed at Stanford University. Y.S. conducted mouse surgeries for one-photon in vivo imaging of interneurons in motor cortex, and maintained the Htr3a-cre mouse line. C.D. conducted mouse surgeries and provided guidance for in vivo imaging of CA1 neurons in Hippocampus. J.Z prepared samples, acquired data, and analyzed data for two-photon in vivo imaging in mouse visual cortex. L.C.G. performed mouse surgeries, prepared samples, acquired data, and analyzed data for two-photon in vivo imaging in mouse barrel cortex. R.G.N. performed mouse surgeries for two-photon in vivo imaging in mouse visual cortex. G.T.S. performed characterization of ASAP6 variants in mouse ex vivo experiments and analyzed the data. A.H. performed characterization of ASAP6 variants in mouse ex vivo experiments. D.J. performed characterization of ASAP6 variants. L.X.L. maintained all mouse lines for in vivo experiments performed at Stanford University and prepared perfused brain samples after in vivo experiments. B.R., D.F., N.J., T.R.C., L.G., and J.D. provided supervision, advised on experimental design, and assisted with data interpretation. M.Z.L.conceived of the project, provided supervision, advised on experimental design, assisted with data interpretation, prepared figures, and co-wrote the manuscript. All authors contributed to the writing of the manuscript.

## Notes

### Competing Interest Statement

M.Z.L. is an inventor on a patent for the earlier ASAP1 voltage indicator. The remaining authors declare no competing interests.

### Summary of Updates

Figure 1 has been revised with ASAP6c data New figures 2-4 have been added Supplementary Tables 2-4 have been revised with ASAP6c data

## REFERENCES

1. Loewi, O. Über humorale übertragbarkeit der Herznervenwirkung. I. Mitteilung. Pflügers Archiv für die Gesamte Physiologie des Menschen und der Tiere 189, 239–242 (1921).

2. Woodbury, J. W. & Patton, H. D. Electrical activity of single spinal cord elements. Cold Spring Harb Symp Quant Biol 17, 185–188 (1952).

3. Bernstein, J. Untersuchungen zur Thermodynamik der bioelektrischen Ströme. Erster Theil. Pflüger Archiv für die Gesammte Physiologie des Menschen und der Thiere 92, 521–562 (1902).

4. Patiño, M., Rossa, M. A., Lagos, W. N., Patne, N. S. & Callaway, E. M. Transcriptomic cell-type specificity of local cortical circuits. Neuron 112, 3851–3866.e4 (2024).

5. Wang, X. J. & Buzsáki, G. Gamma oscillation by synaptic inhibition in a hippocampal interneuronal network model. J. Neurosci. 16, 6402–6413 (1996).

6. Sohal, V. S., Zhang, F., Yizhar, O. & Deisseroth, K. Parvalbumin neurons and gamma rhythms enhance cortical circuit performance. Nature 459, 698–702 (2009).

7. Womelsdorf, T. et al. Modulation of neuronal interactions through neuronal synchronization. Science 316, 1609–1612 (2007).

8. Vancura, B., Geiller, T., Grosmark, A., Zhao, V. & Losonczy, A. Inhibitory control of sharp-wave ripple duration during learning in hippocampal recurrent networks. Nat Neurosci 26, 788–797 (2023).

9. Klausberger, T. et al. Brain-state- and cell-type-specific firing of hippocampal interneurons in vivo. Nature 421, 844–848 (2003).

10. Markram, H., Lübke, J., Frotscher, M. & Sakmann, B. Regulation of synaptic efficacy by coincidence of postsynaptic APs and EPSPs. Science 275, 213–215 (1997).

11. Bi, G. Q. & Poo, M. M. Synaptic modifications in cultured hippocampal neurons: dependence on spike timing, synaptic strength, and postsynaptic cell type. J. Neurosci. 18, 10464–10472 (1998).

12. Song, S., Miller, K. D. & Abbott, L. F. Competitive Hebbian learning through spike-timing-dependent synaptic plasticity. Nat Neurosci 3, 919–926 (2000).

13. Villette, V. et al. Ultrafast Two-Photon Imaging of a High-Gain Voltage Indicator in Awake Behaving Mice. Cell 179, 1590–1608.e23 (2019).

14. Zhang, Y. et al. Fast and sensitive GCaMP calcium indicators for imaging neural populations. Nature 615, 884–891 (2023).

15. e, S. W. et al. A positively tuned voltage indicator for extended electrical recordings in the brain. Nat Methods 20, 1104–1113 (2023).

16. Hao, Y. A. et al. A fast and responsive voltage indicator with enhanced sensitivity for unitary synaptic events. Neuron 112, 3680–3696.e8 (2024).

17. Chamberland, S. et al. Fast two-photon imaging of subcellular voltage dynamics in neuronal tissue with genetically encoded indicators. Elife 6, e25690 (2017).

18. Horn, R. Uncooperative voltage sensors. J Gen Physiol 133, 463–466 (2009).

19. Daigle, T. L. et al. A Suite of Transgenic Driver and Reporter Mouse Lines with Enhanced Brain-Cell-Type Targeting and Functionality. Cell 174, 465–480.e22 (2018).

20. Brinks, D., Klein, A. J. & Cohen, A. E. Two-Photon Lifetime Imaging of Voltage Indicating Proteins as a Probe of Absolute Membrane Voltage. Biophys. J. 109, 914–921 (2015).

21. Wu, J. et al. Kilohertz two-photon fluorescence microscopy imaging of neural activity in vivo. Nat Methods 17, 287–290 (2020).

22. Zhong, J. et al. FACED 2.0 enables large-scale voltage and calcium imaging in vivo. Nat Methods (2025).

23. Lam, A. J. et al. Improving FRET dynamic range with bright green and red fluorescent proteins. Nat Methods 9, 1005–1012 (2012).

24. Lee, S., Hjerling-Leffler, J., Zagha, E., Fishell, G. & Rudy, B. The largest group of superficial neocortical GABAergic interneurons expresses ionotropic serotonin receptors. J. Neurosci. 30, 16796–16808 (2010).

25. Vucurovic, K. et al. Serotonin 3A receptor subtype as an early and protracted marker of cortical interneuron subpopulations. Cereb Cortex 20, 2333–2347 (2010).

26. Dinc, F. et al. Fast, scalable, and statistically robust cell extraction from large-scale neural calcium imaging datasets. bioRxiv 40 (2021).

27. Kannan, M. et al. Dual-polarity voltage imaging of the concurrent dynamics of multiple neuron types. Science 378, eabm8797 (2022).

28. Rupprecht, P., Rózsa, M., Fang, X., Svoboda, K. & Helmchen, F. Spike inference from calcium imaging data acquired with GCaMP8 indicators. bioRxiv 1 (2025).

29. Inoue, M. et al. Rational Engineering of XCaMPs, a Multicolor GECI Suite for In Vivo Imaging of Complex Brain Circuit Dynamics. Cell 177, 1346–1360.e24 (2019).

30. Brooks, F. P. et al. Photophysics-informed two-photon voltage imaging using FRET-opsin voltage indicators. Sci Adv 11, eadp5763 (2025).

31. Grimm, C. et al. Two-photon voltage imaging with rhodopsin-based sensors. Neuron 114, 1198–1209.e7 (2026).

32. Podgorski, K. & Ranganathan, G. Brain heating induced by near-infrared lasers during multiphoton microscopy. J. Neurophysiol. 116, 1012–1023 (2016).

33. Baker, C. A., Elyada, Y. M., Parra, A. & Bolton, M. M. Cellular resolution circuit mapping with temporal-focused excitation of soma-targeted channelrhodopsin. Elife 5, e14193 (2016).

34. Wu, Y. et al. Fast analysis and engineering of protein function by microbe-independent deep assembly and screening. Mol Syst Biol (2026).

35. Pnevmatikakis, E. A. & Giovannucci, A. NoRMCorre: An online algorithm for piecewise rigid motion correction of calcium imaging data. J Neurosci Methods 291, 83–94 (2017).

36. Sun, W., Tan, Z., Mensh, B. D. & Ji, N. Thalamus provides layer 4 of primary visual cortex with orientation- and direction-tuned inputs. Nat Neurosci 19, 308–315 (2016).

37. Brainard, D. H. The Psychophysics Toolbox. Spat Vis 10, 433–436 (1997).

38. Stringer, C. & Pachitariu, M. Cellpose3: one-click image restoration for improved cellular segmentation. Nat Methods 22, 592–599 (2025).

39. Hwang, F. J. et al. Motor learning selectively strengthens cortical and striatal synapses of motor engram neurons. Neuron 110, 2790–2801.e5 (2022).

40. Cai, C. et al. VolPy: Automated and scalable analysis pipelines for voltage imaging datasets. PLoS Comput Biol 17, e1008806 (2021).

41. Drew, P. J. & Feldman, D. E. Intrinsic signal imaging of deprivation-induced contraction of whisker representations in rat somatosensory cortex. Cereb Cortex 19, 331–348 (2009).

42. Fan, J. L. et al. High-speed volumetric two-photon fluorescence imaging of neurovascular dynamics. Nat Commun 11, 6020 (2020).

43. Deneux, T. et al. Accurate spike estimation from noisy calcium signals for ultrafast three-dimensional imaging of large neuronal populations in vivo. Nat Commun 7, 12190 (2016).

