## Supplementary Materials for "Improving positively tuned voltage indicators for faster kinetics and higher contrast"

**Affiliations**
[1]Department of Neurobiology, Stanford University, Stanford, CA, USA
[2]Department of Bioengineering, Stanford University, Stanford, CA, USA
[3]Department of Neurosurgery, First Affiliated Hospital of Sun Yat-sen University, Guangzhou, Guangdong, China
[4]Department of Neurosurgery, Stanford University, Stanford, CA 94305, USA
[5]Department of Neuroscience, University of California, Berkeley, CA, USA
[6]Institute of Molecular and Clinical Ophthalmology Basel, Basel, Switzerland
[7]Department of Ophthalmology, University of Basel, Basel, Switzerland
[8]Department of Physics, University of California, Berkeley, CA, USA
[9]Helen Wills Neuroscience Institute, University of California Berkeley, Berkeley, CA, USA
[10]Molecular Biophysics and Integrated Bioimaging Division, Lawrence Berkeley National Laboratory, Berkeley, CA, USA
[11]Wu-Tsai Neuroscience Institute, Stanford University, Stanford, CA, USA
[12]Chan-Zuckerberg BioHub, San Francisco, CA, USA
[13]Howard Hughes Medical Institute, Stanford University School of Medicine, Stanford, CA, USA
[14]Equal contribution

### Supplementary Data

| | $V_{half}$ (mV) [a] | Per-molecule resting brightness ($F_0/F_{max}$) | Per-molecule brightness change | | Relative brightness change | | Per-molecule SNR, 1 AP ($\Delta F_{AP}/F_0^{1/2}$) | Kinetic index ($\Delta F_{AP}/\Delta F_{100}$) |
|---|---|---|---|---|---|---|---|---|
| | | | 100-mV step ($\Delta F_{100}/F_{max}$) | 1 AP ($\Delta F_{AP}/F_{max}$) | 100-mV step ($\Delta F_{100}/F_0$, %) | 1 AP ($\Delta F_{AP}/F_0$, %) [b] | | |
| ASAP4.0 | 40.41 | 0.24 | 0.30 | 0.06 | 127 | 25 | 0.12 | 0.20 |
| ASAP4.0 T206H (ASAP4.1) | -10.85 | 0.44 | 0.43 | 0.11 | 96 | 25 | 0.17 | 0.26 |
| ASAP4b | 3.24 | 0.28 | 0.47 | 0.09 | 168 | 31 | 0.16 | 0.18 |
| ASAP4e | 35.82 | 0.17 | 0.36 | 0.06 | 212 | 33 | 0.14 | 0.16 |
| ASAP4.0 F148P N149V | -16.65 | 0.42 | 0.41 | 0.10 | 97 | 24 | 0.16 | 0.25 |
| ASAP4.0 F148A N149V T206H | 35.52 | 0.22 | 0.30 | 0.05 | 133 | 23 | 0.11 | 0.17 |
| ASAP4.0 F148D N149V T206H | -18.99 | 0.45 | 0.36 | 0.10 | 81 | 22 | 0.15 | 0.27 |
| ASAP4.0 F148I N149V T206H | 33.57 | 0.16 | 0.37 | 0.06 | 231 | 35 | 0.14 | 0.15 |
| ASAP4.0 F148L N149V T206H | 16.2 | 0.24 | 0.42 | 0.07 | 175 | 28 | 0.14 | 0.16 |
| **ASAP4.0 F148F N149K T206H** | -33.38 | 0.49 | 0.40 | 0.16 | 80 | 33 | **0.23** | 0.41 |
| ASAP4.0 G151K | 22.04 | 0.20 | 0.37 | 0.07 | 185 | 33 | 0.15 | 0.18 |
| ASAP4.0 G151K T206A | -13.47 | 0.32 | 0.47 | 0.10 | 145 | 32 | 0.18 | 0.22 |
| ASAP4.0 F148F N149P G151K T206A | -16.87 | 0.32 | 0.50 | 0.07 | 154 | 21 | 0.12 | 0.14 |
| ASAP4.0 F148F N149D G151K T206A | -41.4 | 0.48 | 0.43 | 0.15 | 90 | 31 | 0.21 | 0.34 |
| **ASAP4.0 F148F N149A G151K T206A** | -48.32 | 0.49 | 0.44 | 0.15 | 89 | 31 | **0.22** | 0.35 |

**Supplementary Table 1.** Summary of ASAP4.1 and ASAP4.0 G151K T206A engineering**.** Note that $F_0$ refers to fluorescence at rest, which is at –70 mV in absolute transmembrane voltage, and $\Delta F_{100}$ refers to fluorescence change upon a 100-mV step, which reaches +30 mV in absolute membrane voltage. [a]Midpoint of F-V curve fit to Boltzmann function. [b]$\Delta F/F_0$ fluorescence change per AP waveform (FWHM = 3.5 ms, amplitude = 100 mV) recorded in whole-cell voltage clamp mode in HEK293A cells at room temperature.

| | $V_{half}$ (mV) [a] | Per-molecule resting brightness ($F_0/F_{max}$) | Per-molecule brightness change | | Relative brightness change | | Per-molecule SNR, 1 AP ($\Delta F_{AP}/F_0^{1/2}$) | Kinetic index ($\Delta F_{AP}/\Delta F_{100}$) |
|---|---|---|---|---|---|---|---|---|
| | | | 100-mV step ($\Delta F_{100}/F_{max}$) | 1 AP ($\Delta F_{AP}/F_{max}$) | 100-mV step ($\Delta F_{100}/F_0$, %) | 1 AP ($\Delta F_{AP}/F_0$, %) [b] | | |
| ASAP6.1 | -19.0 | 0.32 | 0.54 | 0.15 | 164 | 50 | 0.26 | 0.31 |
| ASAP6.2 | -42.0 | 0.49 | 0.45 | 0.15 | 92 | 34 | 0.22 | 0.36 |
| ASAP6.3 | 11.8 | 0.21 | 0.46 | 0.13 | 222 | 70 | 0.29 | 0.31 |

**Supplementary Table 2.** Summary of ASAP6.1, 6.2, and 6.3 engineering**.** Note that $F_0$ refers to fluorescence at rest, which is at –70 mV in absolute transmembrane voltage, and $\Delta F_{100}$ refers to fluorescence change upon a 100-mV step, which reaches +30 mV in absolute membrane voltage. [a]Midpoint of *F-V* curve fit to Boltzmann function. [b]$\Delta F/F_0$ fluorescence change per AP waveform (FWHM = 2.0 ms, amplitude = 100 mV) recorded in whole-cell voltage clamp mode in HEK293A cells at room temperature.

| | ASAP6.1 | ASAP6.2 (6b) | ASAP6.3 (6c) |
|---|---|---|---|
| $\tau_{on}$ (–70 to +30 mV, ms) | 3.99 ± 0.20 | 3.74 ± 0.26 | 3.93 ± 0.15 |
| $\tau_{off}$ (+30 to –70 mV, ms) | 4.89 ± 0.11 | 6.16 ± 0.13 | 4.41 ± 0.16 |

**Supplementary Table 3.** Kinetics measured at room-temperature in HEK293A cells imaged at 800Hz. N = 4, 4, and 7 cells for ASAP6.1, ASAP6.2, and ASAP6.3, respectively.

| | Relative per-neuron brightness | Single-AP \|ΔF/F\| | Predicted relative SNR | Measured relative SNR |
|---|---|---|---|---|
| ASAP4e-Kv | 1.0 ± 0.1 (14) | 32 ± 4 % (5) | 1.0 | 1.0 ± 0.1 (5) |
| ASAP5-Kv | 1.5 ± 0.1 (21) | 34 ± 1 % (10)* | 1.3 | ND |
| ASAP6.1-Kv | ND | 48 ± 9 % (3) | ND | 1.3 ± 0.1 (3) |
| ASAP6b-Kv | 1.5 ± 0.1 (18) | 62 ± 3 % (6) | 2.3 | 1.9 ± 0.1 (6) |
| ASAP6c-Kv | 1.0 ± 0.1 (17) | 98 ± 9 % (5) | 2.9 | 2.7 ± 0.2 (5) |
| pAce-Kv | 28 ± 2 (20) | 24 % * | 3.8 | ND |
| Positron525-Kv | 3.5 ± 0.2 (19) | 14 % * | 0.8 | ND |

**Supplementary Table 4.** Per-neuron measured and estimated SNRs for an AP. Per-neuron relative brightness measured in primary rat cortical neurons transiently transfected with each GEVI. Single AP *ΔF/F* and measured relative SNRs are from the neuron patch results shown in Fig. 1h, otherwise noted. Predicted relative SNRs are based on square root of per-neuron brightness and *ΔF/F* for an AP. The brightness and SNRs were normalized to ASAP4e-Kv. Values are mean ± SEM with number of neurons in parentheses. * Values obtained from original literature.

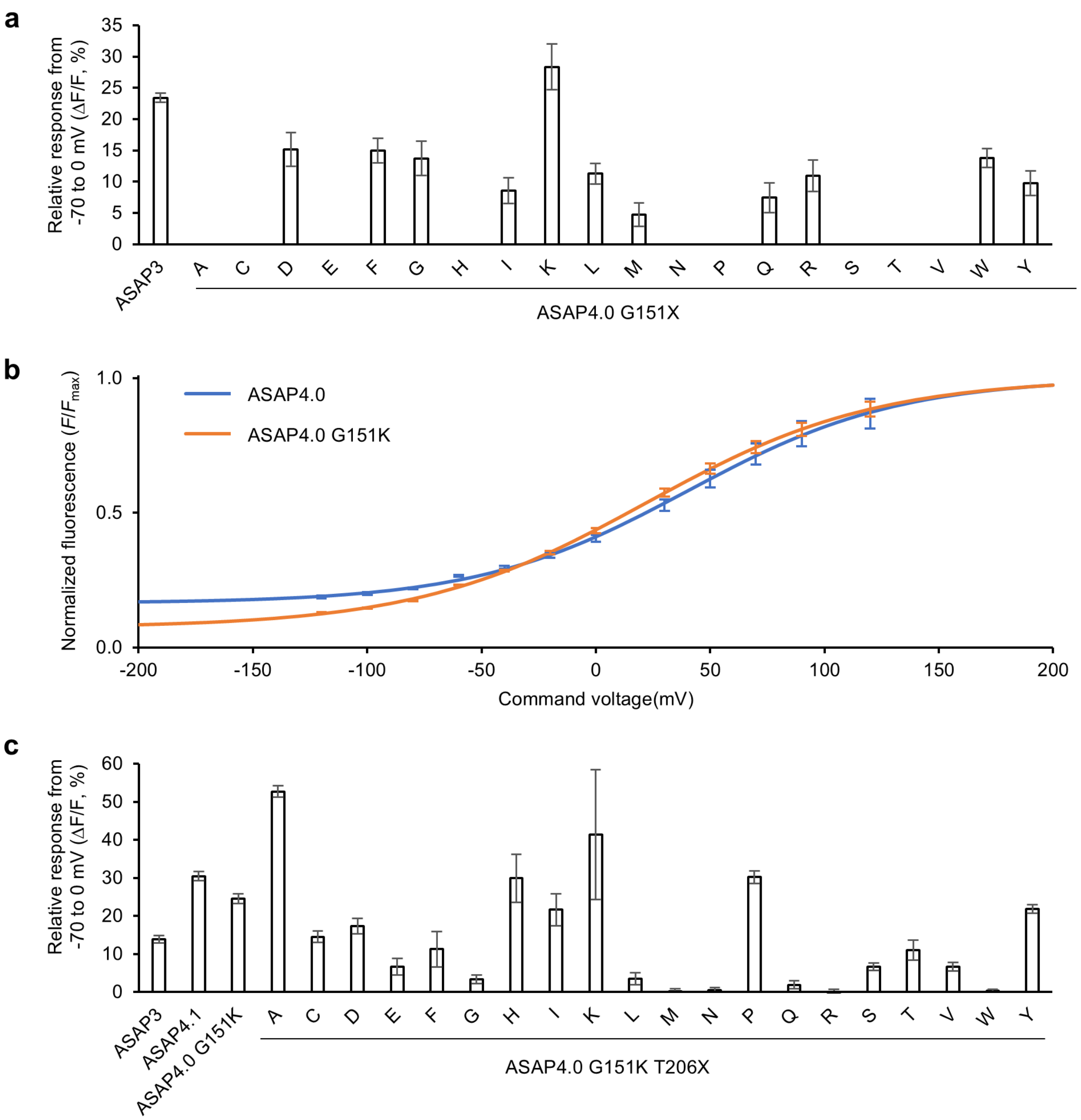


**Supplementary Figure 1.** Screening G151X and T206X of ASAP4.0 yielded ASAP4.0 G151K T206A. **(a)** Peak $\Delta F/F$ responses upon electroporation of HEK293-Kir2.1 cells transfected with PCR fragment library for screening ASAP4.0 D150 G151X variants. ASAP4.0 D150 G151K showed the largest response. **(b)** Resulting F–V curves of ASAP4.0 and ASAP4.0 G151K normalized to $F_{max}$ values. N = 5 (ASAP4.0) and 11 (ASAP4.0 G151K). **(c)** Peak $\Delta F/F$ responses upon electroporation of HEK293-Kir2.1 cells transfected with PCR fragment library for screening ASAP4.0 G151K T206X variants resulted in ASAP4.0 G151K T206A. Error bars are SEM. ASAP3 response was flipped in polarity for display.

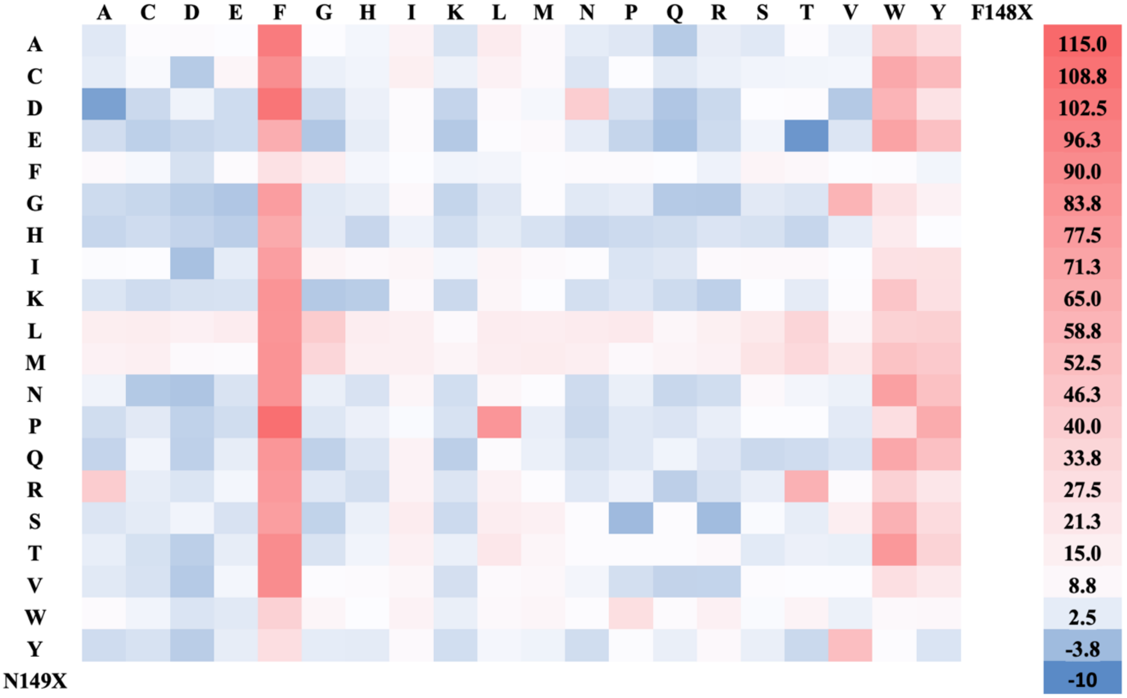


**Supplementary Figure 2.** Screening result of a double-site saturation mutagenesis library of ASAP4.0 G151K T206A F148X N149X. The blue-to-red color scale represents the Δ*F*/*F* (%) response normalized to ASAP4.0 G151K T206A upon electroporation of HEK293-Kir2.1 cells experiencing approximately -70mV to 0mV membrane depolarization.

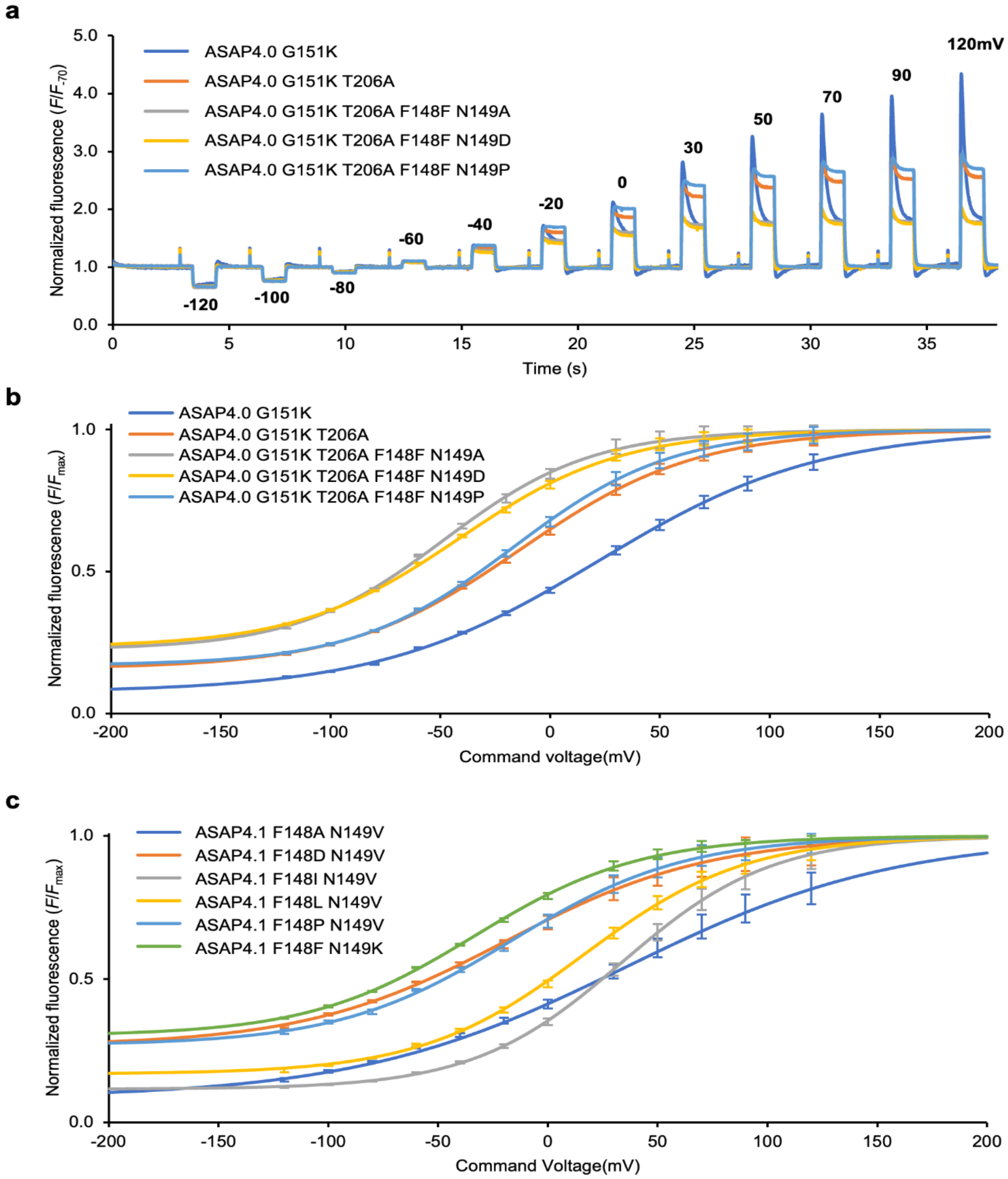


**Supplementary Figure 3.** Fluorescence responses under voltage-clamping of selected variants from ASAP4.0 G151K T206A F148X N149X and ASAP4.1 F148X N149X screening. **(a)** Representative responses of HEK293A cells expressing ASAP4.0 G151K and four variants of ASAP4.0 G151K T206A F148X N149X. The cells were patched in whole-cell voltage clamp mode with command voltage steps. **(b)** Resulting F–V curves of (a) normalized to $F_{max}$ values. N = 11 (ASAP4.0 G151K), 5 (ASAP4.0 G151K T206A), 5 (ASAP4.0 G151K T206A F148F N149A), 5 (ASAP4.0 G151K T206A F148F N149D), and 6 (ASAP4.0 G151K T206A F148F N149P). **(c)** F–V curves of six variants of ASAP4.1 F148X N149X normalized to $F_{max}$ values. N = 6 (ASAP4.1 F148A N149V), 6 (ASAP4.1 F148D N149V), 3 (ASAP4.1 F148I N149V), 3 (ASAP4.1 F148L N149V), 6 (ASAP4.1 F148P N149V), and 6 (ASAP4.1 F148F N149K). Error bars are SEM.

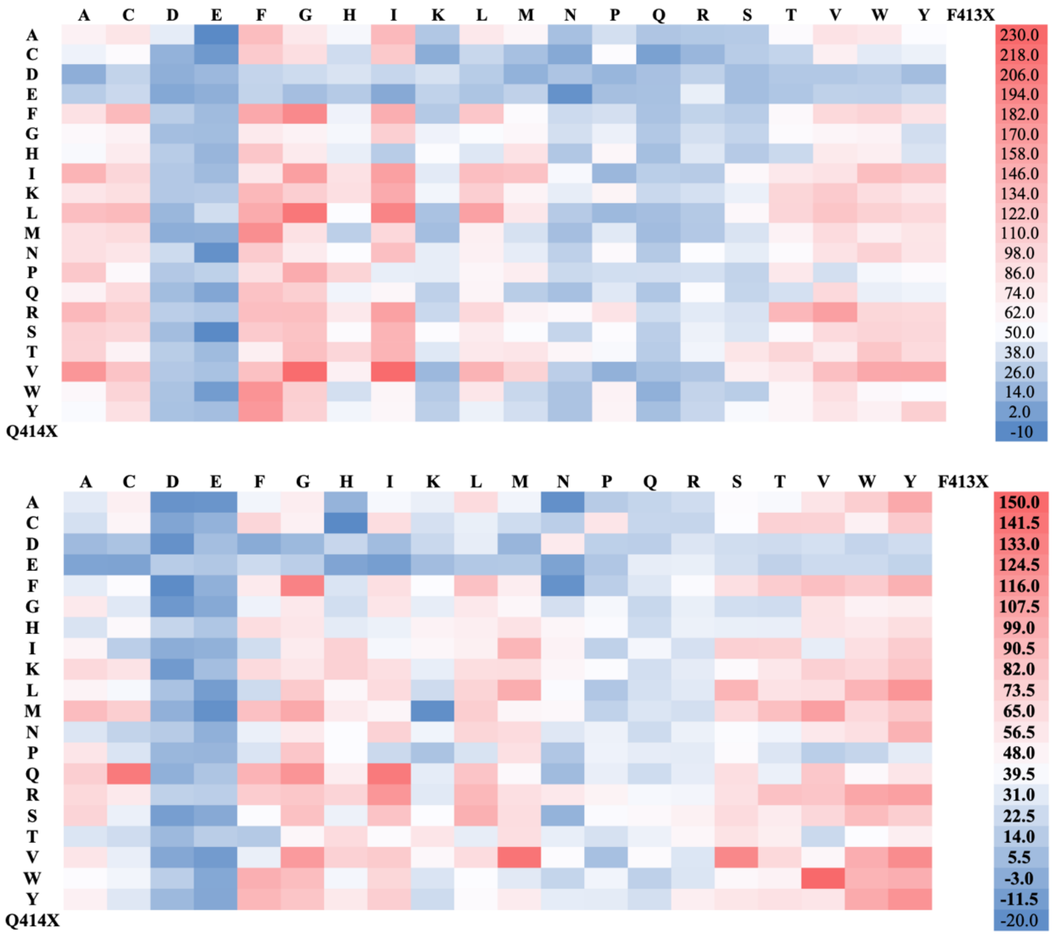


**Supplementary Figure 4.** Screening results of double-site saturation mutagenesis library of ASAP4.0 N149K T206H F413X Q414X (upper panel) and ASAP4.0 N149A G151K T206A F413X Q414X (lower panel) expressed in HEK293-Kir2.1 cells. The blue-to-red color scale represents the Δ*F*/*F* (%) response normalized to ASAP4.0 T206H (top) and ASAP4.0 N149A G151K T206A (bottom) upon electroporation of HEK293-Kir2.1 cells experiencing approximately -70mV to 0mV membrane depolarization.

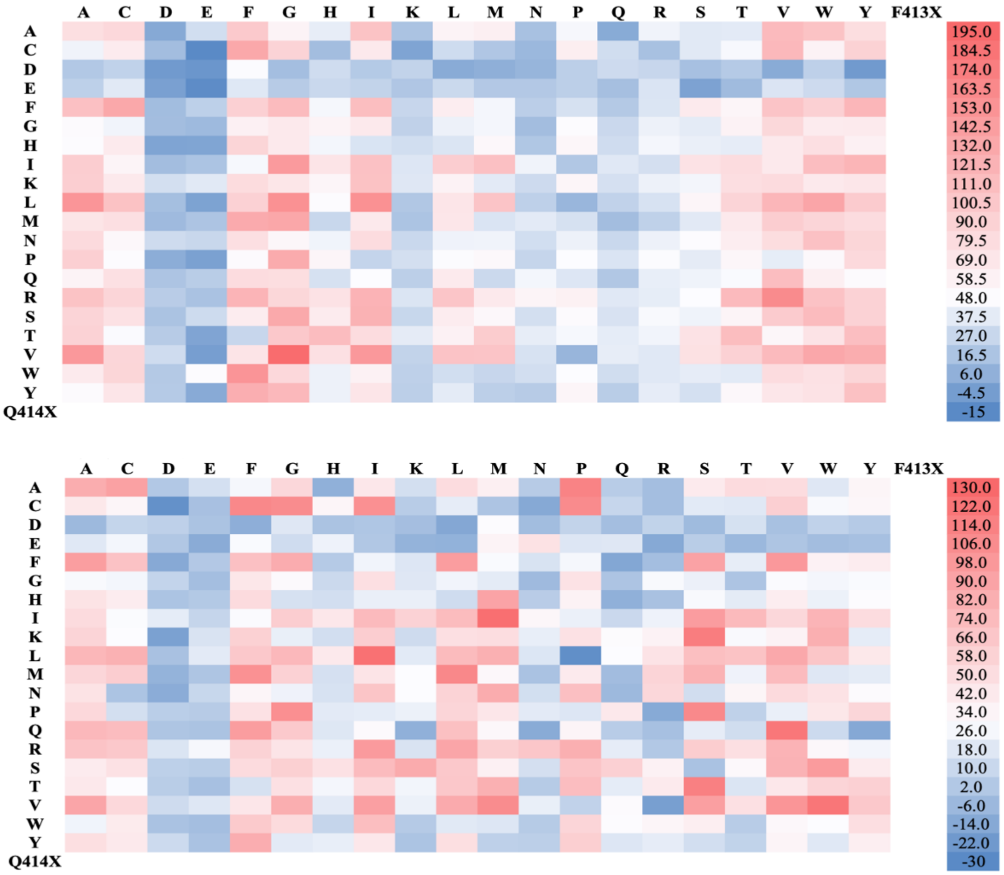


**Supplementary Figure 5.** Screening results of double-site saturation mutagenesis library of ASAP4.0 T206H F413X Q414X (upper panel) and ASAP4.0 G151K T206A F413X Q414X (lower panel) expressing HEK293-Kir2.1 cells. The blue-to-red color scale represents the Δ*F*/*F* (%) response normalized to ASAP4.0 T206H (top) and ASAP4.0 G151K T206A (bottom) upon electroporation of HEK293-Kir2.1 cells experiencing approximately -70mV to 0mV membrane depolarization.

**Supplementary Note**

We initially concentrated on a series of variants with D150X G151X mutations relative to ASAP4.0, as these sites seemed to tolerate multiple amino acids in the screening that led to ASAP4.0[15]. We found ASAP4.0 G151K to have a better responsivity in the electroporation screening and upon voltage-clamp testing **(Supplementary Fig. 1a-1b**), driving a large SNR determinant for AP detection (relative response to a commanded AP waveform multiplied by the square root of basal brightness, **Supplementary Table 1**). We performed screening of T206X mutants on ASAP4.0 G151K, analogous to the mutagenesis of ASAP4.0 that yielded T206H to create ASAP4.1. In ASAP4.0 G151K, T206A stood out with the largest responsivity (**Supplementary Fig. 1c**), in contrast to T206H in ASAP4.1.

Retracing the evolutionary process from ASAP4.1 to ASAP4.2 by yet another step, we used ASAP4.0 G151K T206A as a template for F148X N149X saturated double site mutagenesis. We found multiple high-scoring variants keeping Phe-148 but mutations at Asn-149 which points inward toward the chromophore **(Supplementary Fig. 2-3)**. Upon patch-clamping a subset of higher responders **(Supplementary Fig. 3a-b)**, ASAP4.0 N149A G151K T206A showed the best SNR determinant on simulated APs (**Supplementary Table 1)**. We also patch-clamped ASAP4.1 N149X variants with higher responses in the electroporation screen (**Supplementary Fig. 3c**). We found ASAP4.0 N149K T206H to have brighter baseline fluorescence, larger responsivity, and faster kinetics than the previous ASAP4.2, resulting in a higher SNR determinant for APs (**Supplementary Table 1**). This supplied ASAP4.0 N149K T206H and ASAP4.0 N149A G151K T206A as two leads for the kinetic screen.
